# Rolling Signal-based Ripley’s K: A new algorithm to identify spatial patterns in histological specimens

**DOI:** 10.1101/2020.05.21.109314

**Authors:** Connor P. Healy, Frederick R. Adler, Tara L. Deans

## Abstract

The spatial distribution of cells within a tissue underlies organ function. However, these spatial distributions are often difficult to identify, making it challenging to evaluate how cells establish these patterns during development or how diseases may disrupt these patterns and impair function. To address this, we developed an image analysis tool based on a novel algorithm that identifies spatial patterns within tissues. This analytical tool was used to study the bone marrow, a specialized microenvironment in which spatial patterning of regulatory cells may influence the differentiation and survival of hematopoietic stem cells. Using this algorithm, we discovered clusters of regulatory cells within the bone marrow that suggest an organization of micro-niches, which may form the basis of the hematopoietic stem cell microenvironment. This work provides a new tool for the detection and analysis of tissue morphology that enables identification of spatial patterns within tissues that can lead to a deeper understanding of tissue function, provide clues for early onset of disease, and be used as a tool for studying the impact of pharmaceutics on tissue development and regeneration.

**In Brief:** This work introduces a new statistic to analyze the patterning of cells and physiological features in histological images. This statistic was used on a published set of immunofluorescent images of murine bone to identify novel spatial structures within the bone marrow that may provide new inisghts to the organization of the hematopoietic stem cell microenvironment.

**Highlights:**

- RSRK, a statistical tool for analyzing the spatial distribution of features in histological images, is introduced.
- RSRK incorporates the quantification of signal distribution to identify unique spatial patterns.
- Spatial patterns in hematopoietic stem cell microenvironments are identified.

**Graphical Abstract:** 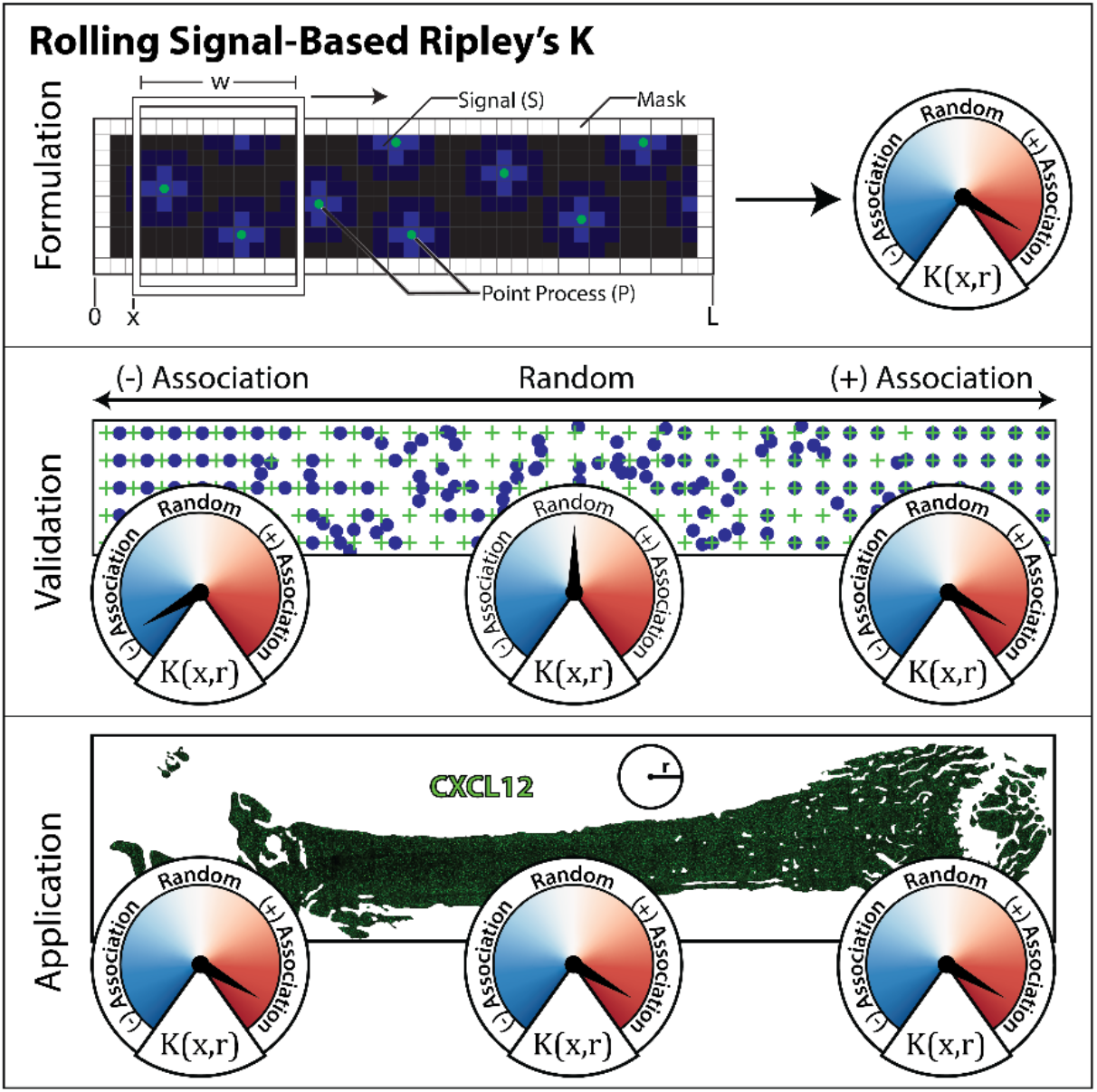

## INTRODUCTION

Spatial patterns among cells can be found in a variety of tissues and organs and often underlie tissue function. For example, photoreceptors form mosaics in the eye that are carefully tuned to permit vision under a variety of lighting conditions (Ahnelt, 1998; Wandell, 1995), the spacing of collecting ducts in the renal medulla is optimized to maximize the filtration capacity of the kidney (Chandhoke et al., 1985), and the macula of the vestibular ear has hair cells that are spaced uniformly amongst support cells (Deans, 2013; Purves et al., 2001). Cellular spatial patterns bridge the gap between single-cell biology and the biology of tissues and systems throughout the body. A better understanding of the mechanisms of how these spatial patterns arise can provide insight into how networks of cells work in concert to contribute to overall tissue function and morphogenesis. Additionally, accurate reproduction of cellular patterning is critical to the successful engineering of mammalian tissues (Javaherian et al., 2013), and first requires the detection, categorization, and analysis of the spatial patterns that may arise in a tissue.

Cell signalling is essential for response to the environmental cues in all multicellular organisms, and coordinates cellular decisions including cell division, apoptosis, differentiation, and migration. Signalling patterns are controlled in part by dividing cells into distinct specialized niches. Malfunctions in cellular signalling that create these niches cause many diseases, including autoimmune and metabolic diseases, cancer, and neurological dysfunctions (Wei et al., 2004). Quantifying the spatial organization of local niches within tissues is thus key for understanding how cells interact and maintain the health of the tissue. The current method for evaluating spatial patterning in tissues is primarily the qualitative analysis of tissue specimens. However, such quantitative analyses are time consuming and prone to intraobserver variability (Bridge et al., 2016).

To more effectively quantify spatial patterning within tissues, we developed a quantitative image analysis tool based on the Ripley’s K statistic (Ripley, 1976). Ripley’s K was initially established for use in the fields of ecology and anthropology to determine whether a spatial feature (e.g., trees in a forest and burial sites) is dispersed, clustered, or randomly distributed throughout an area of interest, and to determine the spatial scale of patterns. Knowing the type and scale of the pattern detected, scientists can infer the cause and mechanisms that produce the pattern (e.g., uniform spatial patterns are a sign of competitive inhibition between members of the same species). While Ripley’s K is a powerful tool for evaluating patterns, it is a global statistic that measures spatial patterns in an entire region and cannot quantify how spatial patterning changes throughout that region (Box 1). Spatially inhomogeneous patterns confound traditional Ripley’s K analysis when patterning in one region masks the opposite patterning in a different region, limiting the application of conventional Ripley’s K to the analysis of spatially homogenous tissue sections (Box 1C). Additionally, Ripley’s K can only analyze point-like patterns and cannot quantify the spatial patterning of point-like features relative to linear and area features (Box 1D). A method that assesses how spatial organization varies throughout an area would overcome these limitations with traditional Ripley’s K and allow for the spatial patterning of cells and features within tissues to be analyzed and quantified.

### Box 1. Ripley’s K

The Ripley’s K statistic was initially developed for use in the fields of ecology and anthropology to summarize the spatial distribution of features in a homogeneous point process (Dixon, 2002; Ripley, 1976). By comparing the Ripley’s K statistic for an observed point process (*K_OBS_*) to the Ripley’s K statistic for random point processes in the same area (*K_RND_*), researchers can characterize the type of spatial patterns that are present in the observed point process at a given scale *r*. For example, the patterns generated by Ripley’s K can give scientists insights into how humans lived centuries ago. One such study examined the spatial patterns of human remains that were discovered in a quarry. Using Ripley’s K to study the burial patterns of the bones, it was determined that the quarry contained a late Prehistoric ossuary {Dirkmaat, 2015 #1335}.

Ripley’s K has two basic formulations, univariate and bivariate, both of which analyze the spatial distribution of points in a point process. Univariate Ripley’s K analyzes the spatial distribution of points of the same type, such as a single cell type. Bivariate Ripley’s K analyzes the spatial distribution of points of two different types e.g., healthy cells vs. cancerous cells. All forms of Ripley’s K have the form

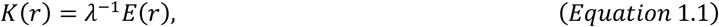

where *E*(*r*) is the expected number of additional points within a radius *r* of a randomly selected point in the *same* point process for univariate Ripley’s K or in a *different* point process for bivariate Ripley’s K. Under complete spatial randomness *E* = *πr*^2^ · *λ* is the density of the point in the area over which the point process is being analyzed. Under both formulations, the Ripley’s K statistic reports the type of spatial patterning that is detected at a given spatial scale. For univariate Ripley’s K, a positive value indicates the point process is clustered while negative value indicates the point process is regularly spaced (Box 1A). For bivariate Ripley’s K positive values indicates a positive association between the two point processes, and negative value indicates a negative association between the two point processes (Box 1B). Additionally, it’s important to note that point processes can and often do exhibit different types of patterns at different scales. The full formulation and implementation of traditional Ripley’s K is reviewed by Dixon (2002).

Traditional Ripley’s K is globally computed and insensitive to shifts in spatial patterning that occur throughout a region. For example, a point process that is clustered on the left and uniform on the right would confound traditional Ripley’s K (Box 1C). Ripley’s K for this particular point process would fail to capture this spatial dependency and conclude that the point process is random at the spatial scales where the two patterns destructively interact. A second limitation of traditional Ripley’s K is its inability to compare the spatial distribution of points with respect to non-point spatial features. For example, the points in the red point process in Box 1D are clustered near the blue area feature. However, since the blue area feature is not a point process, Ripley’s K cannot be used to analyze this interaction.

**Figure B1.**
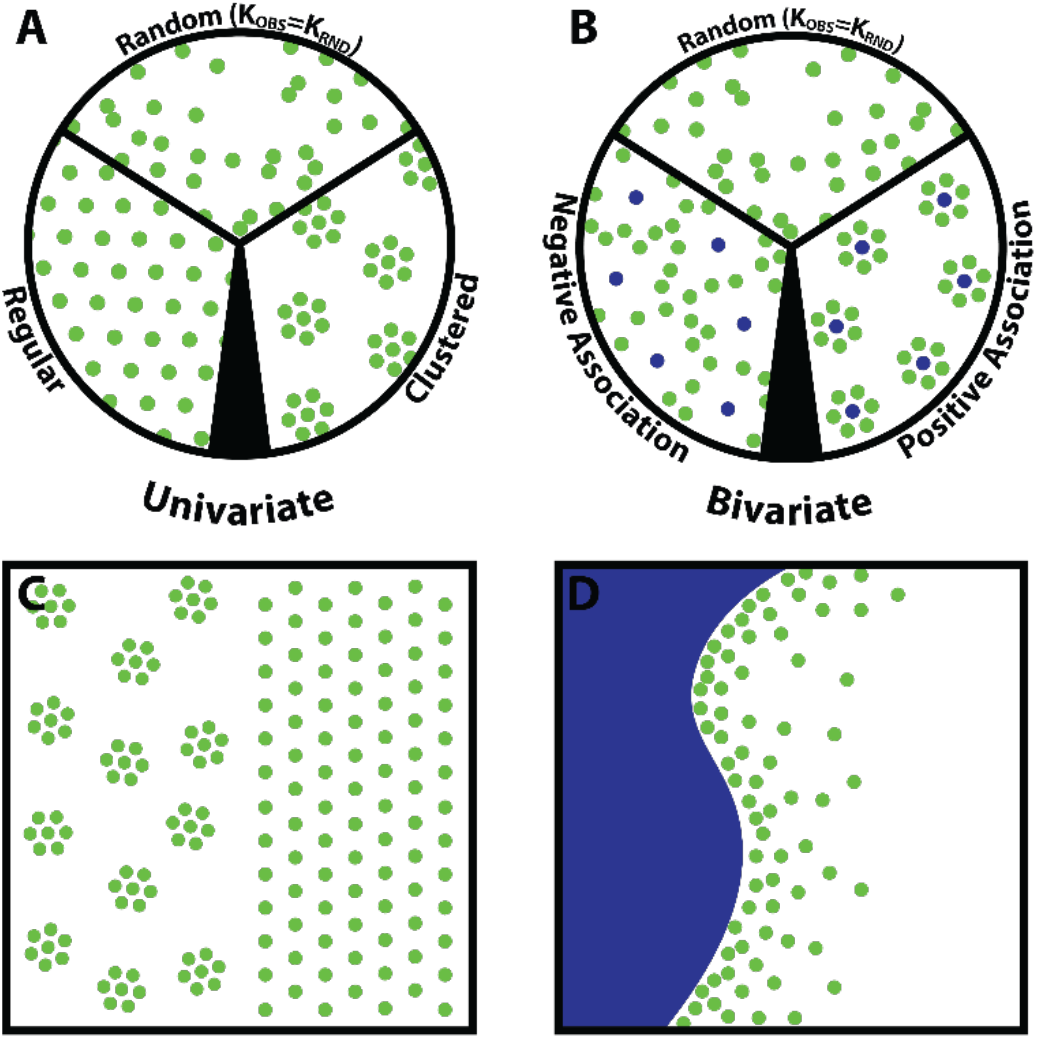
Traditional Ripley’s K. (A) Examples of regular, clustered, and random patterns in a univariate point process. (B) Examples of negatively associated, randomly associated, and positively associated patterns in a bivariate point process. (C-D) Examples of spatial patterns that confound traditional Ripleys K. (C) A univariate point pattern that is equal parts clustered and equal parts uniform. (D) A point process (green) attracted to an area feature (blue).

In this article, we describe and test Rolling Signal-based Ripley’s K (RSRK), a statistic that allows the scanning of biomedical images to identify region-specific spatial patterning between point-like features as well as non-point like features. This approach was applied to study the bone marrow microenvironment, which consists of a diverse population of cells and tissues that work in concert to regulate hematopoietic stem cells (HSCs), the stem cell source of all blood and immune cells (Kumar et al., 2018). HSCs are believed to be maintained within the bone marrow by a cellular niche that is made up, at least in part, by CXCL12+ regulatory cells known to be critical to the maintenance and retention of HSCs (Golan et al., 2013, 2018; Greenbaum et al., 2013; Wright et al., 2002). However, unlike stem cell niches in other tissues, no landmarks or morphological structures have been identify to articulate the HSC niche within the bone marrow (Petrie and Zúñiga-Pflücker, 2007; Wei and Frenette, 2018). To assess whether any of these morphological features may be present in the bone marrow, RSRK was used to search for region-specific spatial patterning among CXCL12+ regulatory cells with respect to themselves, and with respect to other major physiological features of the bone marrow including the endosteum and vasculature. Using RSRK the spatial arrangement of CXCL12+ cells within the bone marrow was measured and it was discovered that CXCL12+ cells form clusters throughout the bone marrow. Our data suggest that the presence of clustering amongst CXCL12+ cells, along with the critical role that CXCL12+ cells play in HSC regulation, suggests that these cell clusters could be a landmark for, or even form the basis of, the elusive HSC niche.

## RESULTS

### Formulating Rolling Signal-Based Ripley’s K

Rolling signal-based Ripley’s K (RSRK) is a variation of Ripley’s K, a statistic for the analysis of spatial point processes (Ripley, 1976). A point process is a set of coordinates defined in a given area (e.g., the position of cells within a tissue specimen). All forms of Ripley’s K take the form:

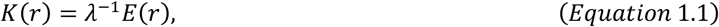

where E(r) is the expected number of additional points within a radius *r* of a randomly selected point in the *same* point process for Univariate Ripley’s K or a *different* point process for Bivariate Ripley’s K. *λ* is the number of points in the area that defines the point process, also known as the density, and its form depends on the type of Ripley’s K analysis being performed (Box 1). *K*(*r*) is the Ripley’s K statistic for a given radius or scale *r* and indicates the type of spatial patterning at that scale; the type of patterning can be random (independent), clustered (positively associated), or regular (negatively associated) (Figure B1 A-B). To simplify the interpretation of the RSRK statistic, most implementations of Ripley’s K compare an observed *K*(*r*) to the *K*(*r*) of a randomly generated point process. An observed *K*(*r*) that is greater than the *K*(*r*) from a randomly generated point process indicates positively associated patterns, e.g. cluster formation and attraction, in the observed point process(es) at spatial scale *r*. Conversely, an observed *K*(*r*) that is less than the *K*(*r*) of a randomly generated point process indicates negatively associated patterns, e.g. regular-spacing, repulsion, and avoidance, in the observed point process(es) at spatial scale *r*.

In biological tissues many spatial patterns arise with respect to spatial features that are not inherently point-like. This prevents analysis of the spatial patterning of point-like features, like cells, with respect to blood vessels, bone, and other major physiological features. RSRK is able to analyze patterning of points with respect to non-point like features by evaluating the intensity of a signal within a distance *r* of a randomly selected point in the point process being analyzed. This signal can be any measurement that is continuous throughout the area being analyzed. In biomedical applications, using a immunofluorescent image as the input signal is particularly useful because a point process generated from one color channel of an immunofluorescent image can be compared to a non-point like feature depicted in a different color channel of the image by using that channel as the *signal* input to RSRK. A point process generated from one channel can also be compared to the signal from that same channel to evaluate self-patterning. To incorporate signal into RSRK, *E*(*r*) is defined as the expected additional signal within a radius *r* of a randomly selected point in a point process, as opposed to the expected number of additional points within a radius *r* of a randomly selected point in a point process (Box 2). This change makes RSRK more input agnostic compared to traditional Ripley’s K approaches that use only point processes as inputs.

#### Box 2. Rolling Signal-based Ripley’s K

Rolling signal-based Ripley’s K (RSRK) is an expansion upon the function described by B.D. Ripley for analysis of the spatial organization of homogeneous point processes (Ripley, 1976). Traditional Ripley’s K computes a summary statistic describing the spatial distribution of a point process with respect to the same or another point process across an entire region. RSRK computes a summary statistic describing the spatial distribution of a point process *P_i_* with respect to a signal *S* (usually a grayscale image) in a series of windows that span the full region. Additionally, RSRK allows for the analysis of spatial distribution in irregular areas by enabling the user to define regions that are (and are not) to be analyzed by using binary masks (**Figure B2A**). Binary masks are simply images that assign a value of 1 to regions to be analyzed and a value of 0 to regions that are not to be analyzed.

In RSRK, the position of the windows with respect to the entire region is denoted by *x* such that *x* ∈ [0, *L* − *w*] where *L* is the length of the full region, *h* is the window height, and w is the window width and is given by

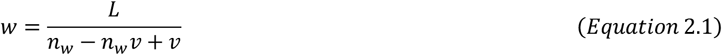

where *n_w_* and *v* are user-defined parameters and correspond to the number of windows spanning the region and desired overlap between windows (expressed as a fraction), respectively. As in traditional Ripley’s K, the window width *w* and height *h* should be defined such that they are at least twice the maximum distance scale *r_max_* that is to be analyzed (Dixon, 2002).

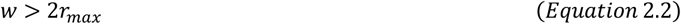

The spacing between window positions Δ*x* is given by

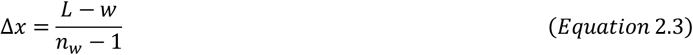

Rolling signal-based Ripley’s K *K*(*x,r*) is given by

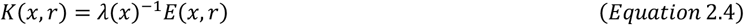

where *λ*(*x*) is the normalization factor for the window with position *x* and is given by

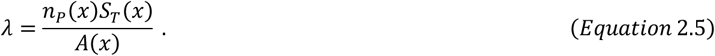

Here *n_P_*(*x*) is the number of points in the window, *S_T_*(*x*) is the total signal in the window with the mask applied (usually the total intensity of all pixels in the window), and *A*(*x*) is the total *unmasked* area in the window (usually the sum of all unmasked pixels).

*E*(*x,r*) is the expected additional signal *S*(*x*) within a radius *r* of a randomly selected point in the point process *P*(*x*) for the window with position *x* and is given by

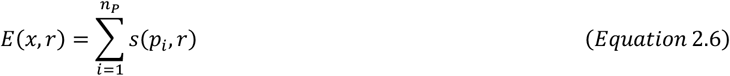

where *s* is a function that gives the total signal intensity in the circle with radius *r* centered at *p_i_* (the *ith* point in *P*(*x*)) (**Figure B2B**).

The null hypothesis for RSRK is that the point process and signal are independent, or randomly placed with respect to each other. To improve the interpretability of *K*(*x,r*) and to test RSRK results against the null hypothesis, the observed RSRK *K_OBS_*(*x,r*) (**Figure B2B**) is compared to the RSRK results of N randomly simulated point processes *K_RND_*(*x,r,n*). The points from the observed point process are set to a random location within the unmasked area of the current window and RSRK is computed. The process is then repeated *N* times (**Figure B2C**). *K_RND_*(*x, r, n*) can then be used to compute significance envelopes with significance levels *α* given by

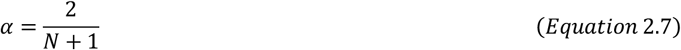

Values of *K_OBS_*(*x, r*) that fall outside the range defined by these significance envelopes are considered significant with a significance level *α*. *K_RND_*(*x,r,n*) can be used to compute an average RSRK for a random point process 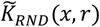 given by

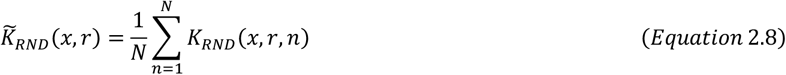

We use 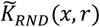 to calculate the fold excess signal within a radius *r* of a random point in the point process *G*(*x, r*)

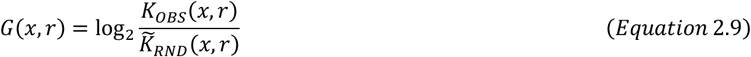

In traditional Ripley’s K, edge effects must be corrected for with an edge correction term (Goreaud and Pelissier, 1999). The normalization used in RSRK automatically adjusts for these edge effects. Furthermore, interpretation of *G*(*x, r*) gives both the type and magnitude of the spatial relationship between the point process and the signal. *G* ≈ 0 indicates that the point process and signal are independent. *G* > 0 demonstrates that there is a positive association (or clustering) between the point process and the signal. *G* < 0 indicates that there is a negative association (or regularity) between the point process and the signal.

It is also possible to pool data from multiple subjects or images to obtain a single combined RSRK value. For *m* ∈ [1, *M*] samples, pooled RSRK (*K_M_*) is given by

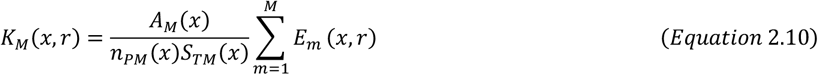

where

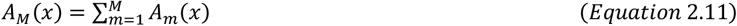

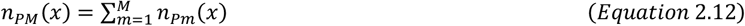

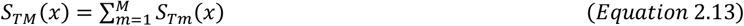

Masks with particularly complex geometry can result artifacts in the RSRK statistics when the scale *r* exceeds the area of the unmasked regions in the image. To address this we define *r_int_* as the maximum interpretable *r* for a masked image. *r_int_* is given by

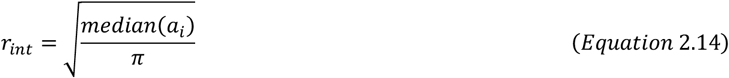

where *a_i_* is the set of areas of all unmasked regions in the image such that *r_int_* is equal to the radius of the circle with area equal to the median of the set of areas. For pooled data, use the median *r_int_* computed for the combined set of areas from all pooled images.

**Figure B2.**
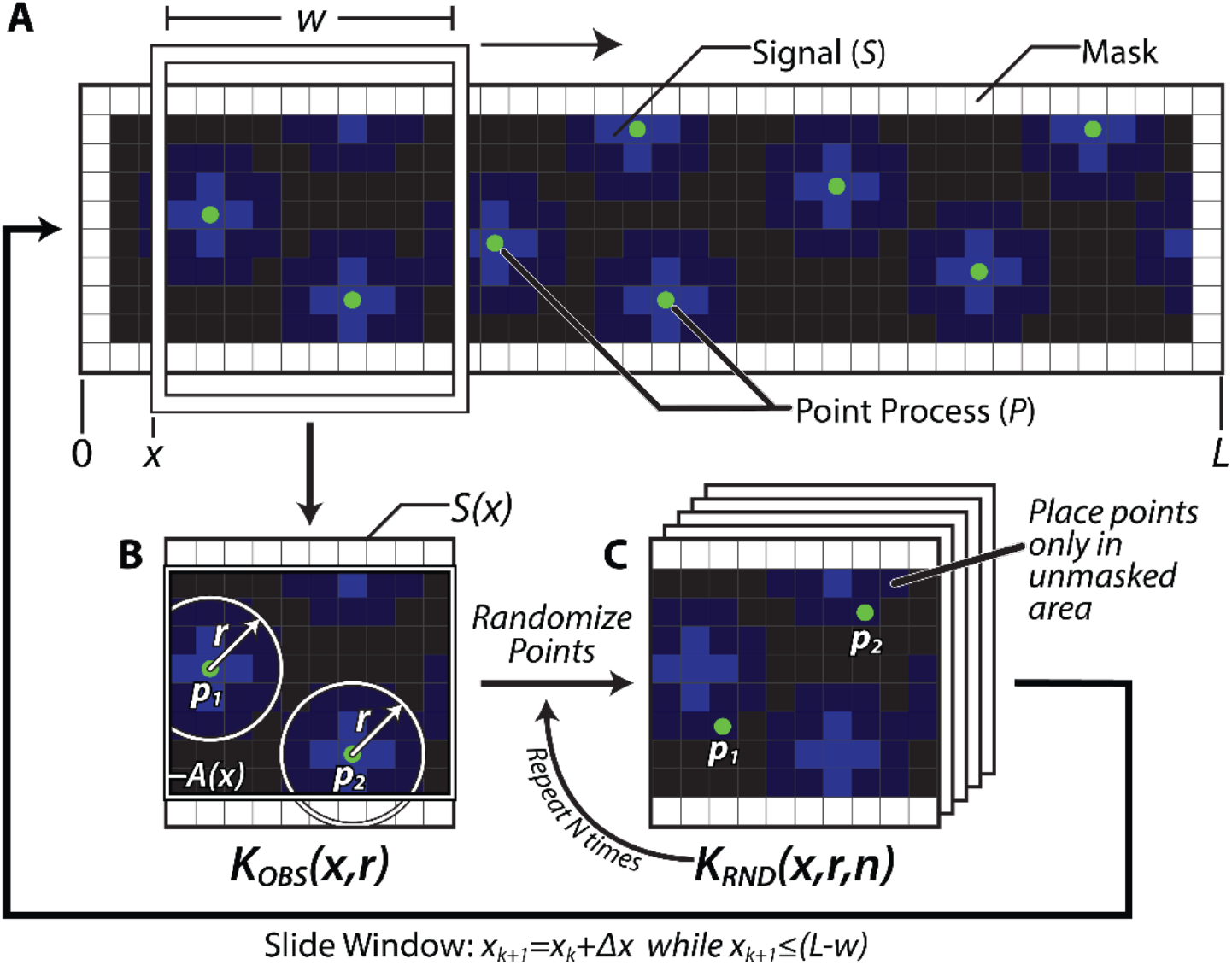
Computation of Rolling Signal-based Ripley’s K. **(A)** Sliding window scheme. Rolling Signal-based Ripley’s K (RSRK) is computed in a series of windows with width *w* across the length *L* of an entire image. The critical inputs for RSRK are the point process *P* (green points) and the Signal S being compared to the point process (blue). An optional mask can be applied to perform RSRK on a region of interest. **(B)** Computation of the observed RSRK statistic (*K_OBS_*). *K_obs_*(*x,r*) is computed for each window with location *x* at the scale *r* using the observed signal *S* and point process *P* as inputs according to **Equation 2.4.** **(C)** Computation of RSRK for a random point process. The points in the observed point process are randomly relocated, and the RSRK statistic is recomputed a total of *N* times to compute the expected RSRK statistic for random spatial organization in the given window 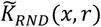 according to *Equation 2*.

To characterize local patterns, a moving window scheme was implemented. RSRK defines a window that scans across an image and computes a RSRK statistic at each position of the window. This addresses the inability of traditional Ripley’s K to characterize spatial patterning throughout a region and enables RSRK to assess changes in spatial patterning that occur across a region. Like traditional Ripley’s K, RSRK also compares the observed RSRK statistic to the RSRK statistic from a randomly generated point process to improve interpretations (Box 2).

### Validating Rolling Signal-Based Ripley’s K

To evaluate RSRK, a set of pre-defined test images was used containing a point process and signal in pre-defined patterns, densities and arrangements. This allowed the evaluation of RSRK to detect different types of patterns, subtle differences in patterns, and to get an overall sense of the robustness of the RSRK statistic, as well as gain an intuition for the interpretation of RSRK results. In the first set of verification tests, the capacity of RSRK to evaluate the spatial arrangement of point processes was characterized with respect to continuous signals. For this verification step, test images were generated of a point process (green) and a signal (blue) arranged in a variety of spatial patterns. Initially, the three primary types of spatial patterns that can exist between a point process and a signal were tested: perfectly aligned (positively associated) with each other, randomly arranged with respect to each other, and perfectly misaligned (negatively associated) with each other (Figure 1A). The RSRK statistic for the perfectly aligned test image indicates significant positive association at scales up to 20 pixels, while the statistic for the perfectly unaligned test image indicates significant negative association at scales up to 15 pixels (Figure 1A). The RSRK statistic for the random test image indicates patterning that is not significantly different from random at all scales (Figure 1A). The aligned and unaligned test results converge to randomness at higher scales because at these scales the point process and signal do not differ significantly from random. These tests confirmed that RSRK can detect the 3 major types of spatial patterning detected by traditional Ripley’s K statistics: positive spatial associations (clustering and attraction), negative spatial associations (regularity and repulsion), and random spatial association (independence).

**Figure 1.**
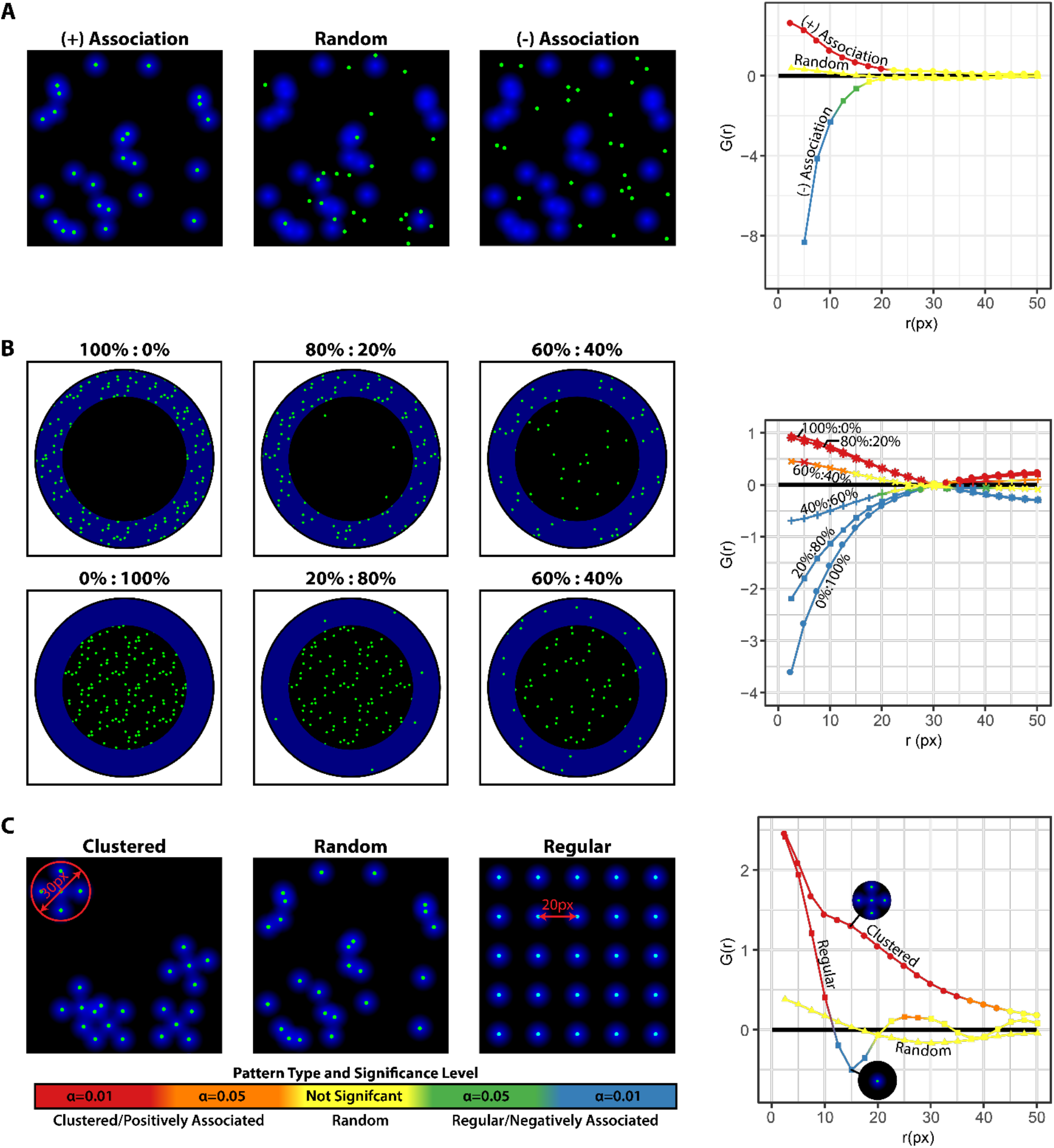
RSRK captures spatial patterning between point processes and signals. A series of test images were designed to test RSRKs sensitivity and ability to detect various types of spatial patterns between signals (blue) and points (green). White regions indicate where a mask was applied to restrict the shape and size of the analysis region. At the right of each set of test images, the RSRK results, for a non-moving scheme with a single window, are provided. The color of each data point indicates the significance and type of spatial patterning detected by RSRK. (A) Testing the 3 possible ideal spatial relationships that can exist between a point and a signal: perfectly associated, entirely random, and perfectly disassociated. Each of these contain the same number of points and total signal but in different spatial arrangements with respect to each other. The radius of the signal circles 7.5 pixels and the total area is 100×100 pixels. (B) Testing the sensitivity of RSRK to shifts in point density. Two regions with equal area are defined in each image. Region A, a ring concentric with the circular mask in which signal is present, and Region B, the circle at the center of Region A in which no signal is present. Point processes at different densities (expressed as percentage of maximum density in the form Region A %: Region B %) were then placed in defined in each region before computing the RSRK statistic for each test image. (C) Additional test images were designed to evaluate the effect that self-expression (small scale association between a point and signal) has on the detection of patterns at larger scales. All points were placed at the center of a 7.5 radius signal circle then the arrangement of these signal point pairs was organized into clustered, randomly positioned, or uniformly spaced patterns. 25 point/signal pairs were used in each test image. Snapshots of the spatial pattern observed at a given scale are included at key points in the RSRK plot.

In the next set of verification tests, we sought to characterize the ability of RSRK to detect positively associated and negatively associated patterns of different strengths. To accomplish this RSRK analysis was performed on a series of test images in which the density of points colocalized with the signal and the density of points not colocalized with the signal were specifically defined. These test images ranged from strong patterns (e.g. 100% of the points are colocalized with the signal) to weaker patterns (e.g. 40% of all points are colocalized with the signal and 60% are not colocalized with the signal) (Figure 1B). The results of RSRK analysis of these test images demonstrated suitable responsiveness to variations in the densities of the points in the different regions. All test images in which more points were colocalized with the signal than were not colocalized with the signal resulted in positive RSRK statistics at scales less than 30 pixels; indicative of a positive spatial association between the point process and signal at those scales (Figure 1B). The opposite result was detected in test images in which the point process was predominantly not colocalized with the signal. All RSRK statistics were negative for these test images at scales less than 30 px (Figure 1B) indicating a negatively associated spatial pattern. The RSRK statistics for all six test images converge to a random pattern at r = 30 px which is roughly the radius of the circle that defines region B because a given point in the point process is equally likely to be in region A as it is to be in region B, negating any patterns that we imposed in our test images. These results demonstrate that RSRK is able to assess the relative strength of different types of spatial patterns.

These verification tests also highlighted a possible artifact in the RSRK results. For all verification images at scales *greater than* 30 pixels, RSRK predicts the opposite type of patterning detected at scales less than 30 pixels because, at these scales, points in Region A start to see less signal than if they were randomly positioned, while the opposite is true for points in Region B (Figure 1B). This artifact of RSRK analysis prompted the definition of the maximum interpretable r (*r_int_*) for a given image/window as equal to radius of the circle with area equal to the largest unmasked region in the image (Equation 2.14).

Thus far, the ability of RSRK to characterize spatial patterns between two different kinds of features (bivariate analysis) has been tested, e.g. cells versus blood vessels. The next step was to characterize the ability of RSRK to characterize spatial patterns between a point process and signal obtained from the *same* feature (univariate analysis). Under this type of analysis the signal and point process are obtained from the same image or channel for composite images. To rule out that RSRK would be unable to detect large scale patterns due to the influence of small-scale self-expression, large scale patterns in the presence of small scale positive association, several test images were prepared in which a point process and signal are perfectly aligned at small scales (similar to what we would expect under a self-expression pattern) but organized into either a clustered (positively associated), random, or regularly-spaced (negatively associated) pattern at larger scales (Figure 1C). Data indicate that the correct type of spatial pattern at larger scales including the small scale positive association between point processes and signal in both the clustered and unvorm test images (Figure 1C). Specifically, the RSRK results for the clustered test image indicated significant positive association (clustering) at scales up to 40 pixels including a small hump at r = 15 px corresponding to the approximate radius of the clusters in the test image (Figure 1C). The RSRK results for the random test image indicated randomness at *all* spatial scales (Figure 1C) and the results for the regular test image indicated positive association at small scales (due to self expression) then negative association (regularity) at scales in the range 12.5 to 17.5 pixels, corresponding roughly to the spacing between the points in the test image, before reverting to near random spatial arrangement at larger scales (Figure 1C). Altogether, these results demonstrate that small-scale self-expression in univariate RSRK analyses does not obscure the detection of spatial patterns at larger scales. These tests also demonstrated that metrics that define different types of spatial patterns (e.g. cluster radius and spacing between uniform points) are visible as distinct features (e.g. peaks, inflection points, and valleys) in the RSRK results.

To evaluate how effectively the rolling-window component of RSRK detects regional variation in spatial patterning, test images of a point process (green crosses) and a signal (blue circles, 30 pixels in radius) were generated and arranged such that they transition from perfectly unaligned (negatively associated) on the far left of the analysis region, to randomly arranged in the center of the analysis region, to perfectly aligned (positively associated) on the far right of the analysis image (Figure 2A). RSRK analysis of this test image accurately predicted a smooth transition between negative association on left (dark blue), to insignificant versus random patterning in the center (cross-hashed regions), to positive association on the right (dark red) (Figure 2B). In order to evaluate more complex regions, e.g. histological specimens, RSRK requires that a mask be defined to restrict the analysis area. To test the robustness of the RSRK to masking, a binary mask to the first test image was applied and the analysis was repeated (Figure 2C). Results show that RSRK correctly predicted the type of the patterning in each region of the image (Figure 2D). Altogether, the number of significant data points in the results from the analysis of the masked image decreased, corresponding to a reduction in the amount of input data.

**Figure 2.**
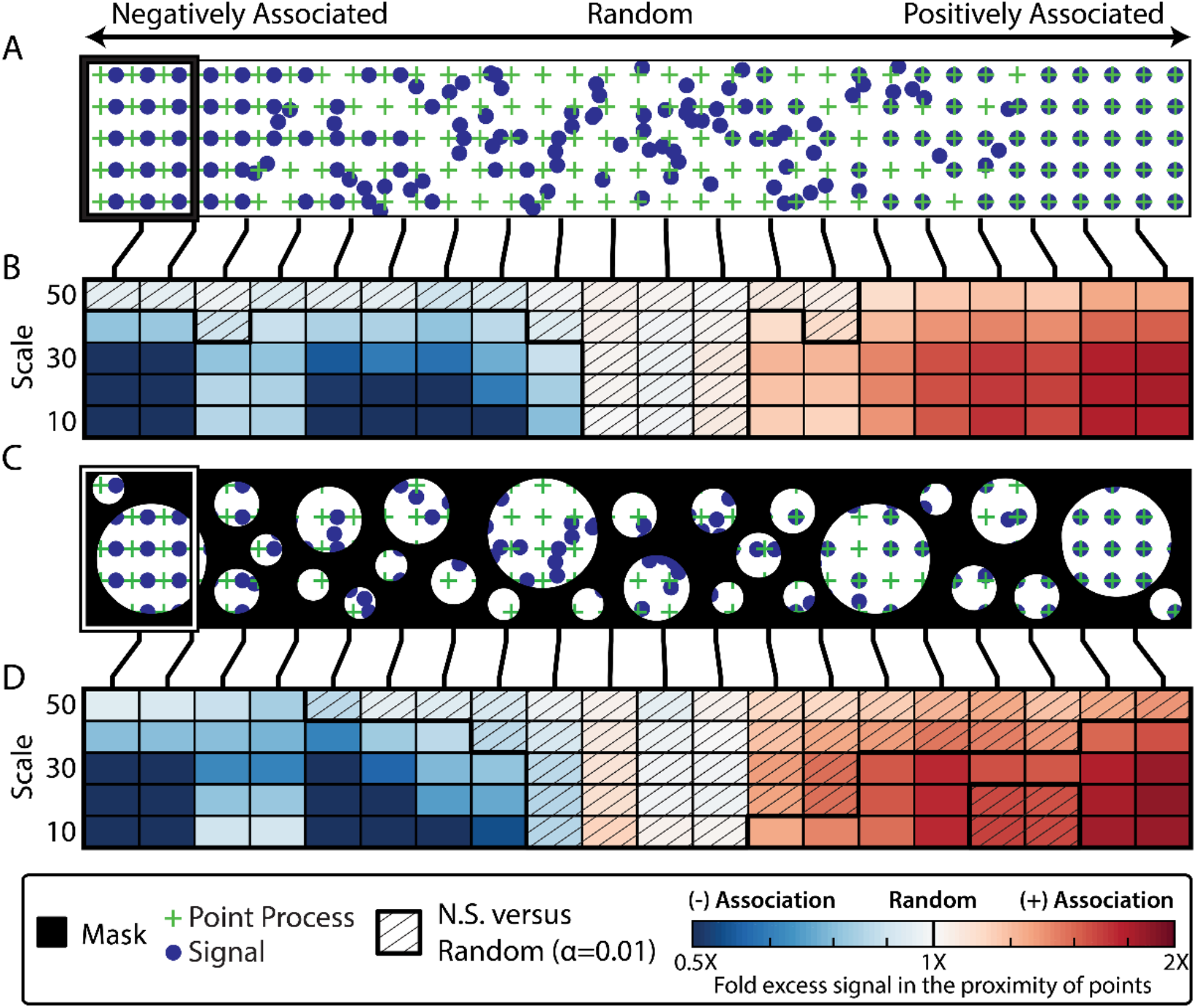
RSRK correctly predicts spatial patterns in masked test images. (A,C) Masked images and point processes used to test RSRK. From left to right, the point process (green) and the signal (blue) transition from perfectly unaligned (negatively associated), to randomly associated, to perfectly aligned (positively associated). The mask applied to the image is shown in black. No mask was applied in A. Example windows are provided on the left of each test image. (B,D) RSRK plots for the images and point processes in (A,C). The hue indicates the type of pattern, and saturation indicates the strength of the pattern. Cross-hatched tiles are not significant (N.S.) versus random. All other tiles are significant versus random. (α=0.01, n=199). Additional Masks are tested in Supplementary Figure S2.

As with prior tests, the rolling window validation test also demonstrated that RSRK correctly identified the scale at which the patterns occurred (Figure 2). The strongest RSRK result for positive association was at *r* = 30*μm*, matching the radius of the blue circles in the signal. Additional testing by making the pattern uniform and the signal transition from positively associated to negatively associated with the signal, and applying different kinds of masks had similar results (Supplementary Figure S2). Altogether, the results of these verification tests provide evidence that RSRK can distinguish between different types of spatial patterns, and delineate between subtle variations in patterning, in a manner that is specific to both the scale at which the patterns occur and the region in which the patterns occur.

### Applying Rolling Signal-Based Ripley’s K to the Bone Marrow

Hematopoesis is a process that takes place in the bone marrow where hematopoietic stem cells (HSCs) produce all blood and immune cells for the body (Kumar et al., 2018). Multiple complex interactions between cells within the bone marrow microenvironment influence HSC self-renewal, proliferation, and the differentiation cues that direct cell fate, as well as the pathophysiology of hematological malignancies (Hoggatt et al.; Kumar et al., 2018). However, studies of the bone marrow have yet to elucidate the interactions and the role of individual cell populations within the HSC niche and how these interactions impact cell fate decisions and/or produce disease. A better understanding of hematopoiesis and the mechanisms that drive hematopoietic disease progression require a clearer picture of the complex HSC regulatory interactions in the bone marrow microenvironment. The spatial patterning of cells and features is anticipated to be fundamental to the function of the bone marrow and has yet to be fully characterized (Boulais and Frenette, 2015; Choi and Harley, 2017; Crane et al., 2017a; Wei and Frenette, 2018); while many extra-medullary hematopoietic organs contain specific spatial structures or landmarks in which hematopoiesis occurs, no such structure has been identified within the bone marrow (Petrie and Zúñiga-Pflücker, 2007; Wei and Frenette, 2018);

To better understand the interactions within the bone marrow that may drive HSC cell fate decisions, the analytical pipeline was applied (semiautomatic image segmentation followed by RSRK analysis) to histological specimens of the bone marrow. The bone marrow as a suitable target for this analysis for various reasons: 1, the bone marrow is a complex organ containing a variety of different cells that work in concert to produce and maintain blood and immune cells via the tight regulation of hematopoietic stem cells; and 2, an extensive database of immunofluorescent images of the bone marrow exists that is ideal for analysis using RSRK (Coutu et al., 2017). Therefore, the RSRK methodology was applied to this database to assess the spatial patterning of a variety of different physiological features and data was pooled from several subjects.

For the analysis, the goal was to capture and analyze the spatial organization of CXCL12+ regulatory cells in the bone marrow with respect to markers for three major physiological regions of the bone marrow: CD31, a marker for vasculature to identify the perivascular region; collagen 1 (ColI), a marker for bone to identify the endosteal region; and the leptin receptor (LepR) to identify cells in the area spanning the other two regions (stromal region). CXCL12+ cells were targeted for RSRK analysis because they are critical to the maintenance and retention of HSCs (Golan et al., 2013, 2018; Greenbaum et al., 2013; Wright et al., 2002), and because evaluation of the spatial arrangement of CXCL12+ cells to the perivascular, endosteal, and stromal markers would provide insights into how each region regulates HSCs (Aiuti et al., 1997; Ara et al., 2003; Crane et al., 2017b; Golan et al., 2013; Greenbaum et al., 2013; Lévesque et al., 2003; Sugiyama et al., 2006; Tzeng et al., 2011; Wei and Frenette, 2018; Wright et al., 2002). Finally, identifying the spatial patterning of CXCL12+ cells within each region of the bone marrow could reveal mechanisms that explain why each region has a different regulatory role with respect to HSCs.

To validate the functionality RSRK analysis of CXCL12+ cells and other features of the bone marrow, additional verification tests were performed in the context of the bone marrow. Initially, the verification test previously discussed (Figure 1B) and masked (Figure 5A) to define the analysis region in the test and to include the the complex geometry of the bone marrow. The test images that were prepared for this verification defined Region A as all areas of the marrow stroma within 90μm of the edge of the endosteal surface (edge of the bone). Region B was defined as all areas within the marrow stroma but not within Region B, effectively the central marrow cavity. As in the previous test, the density of points in each region was varied. Collagen 1 fluorescence (which marks the bone) was used as the signal in this test, resulting in the point process and the signal never colocalizing. In this context a positive association would indicate that a given point is likely to be more dense near the bone (in Region A) than distant from the bone (in Region B). A negative association indicates the opposite (Figure 3).

**Figure 3.**
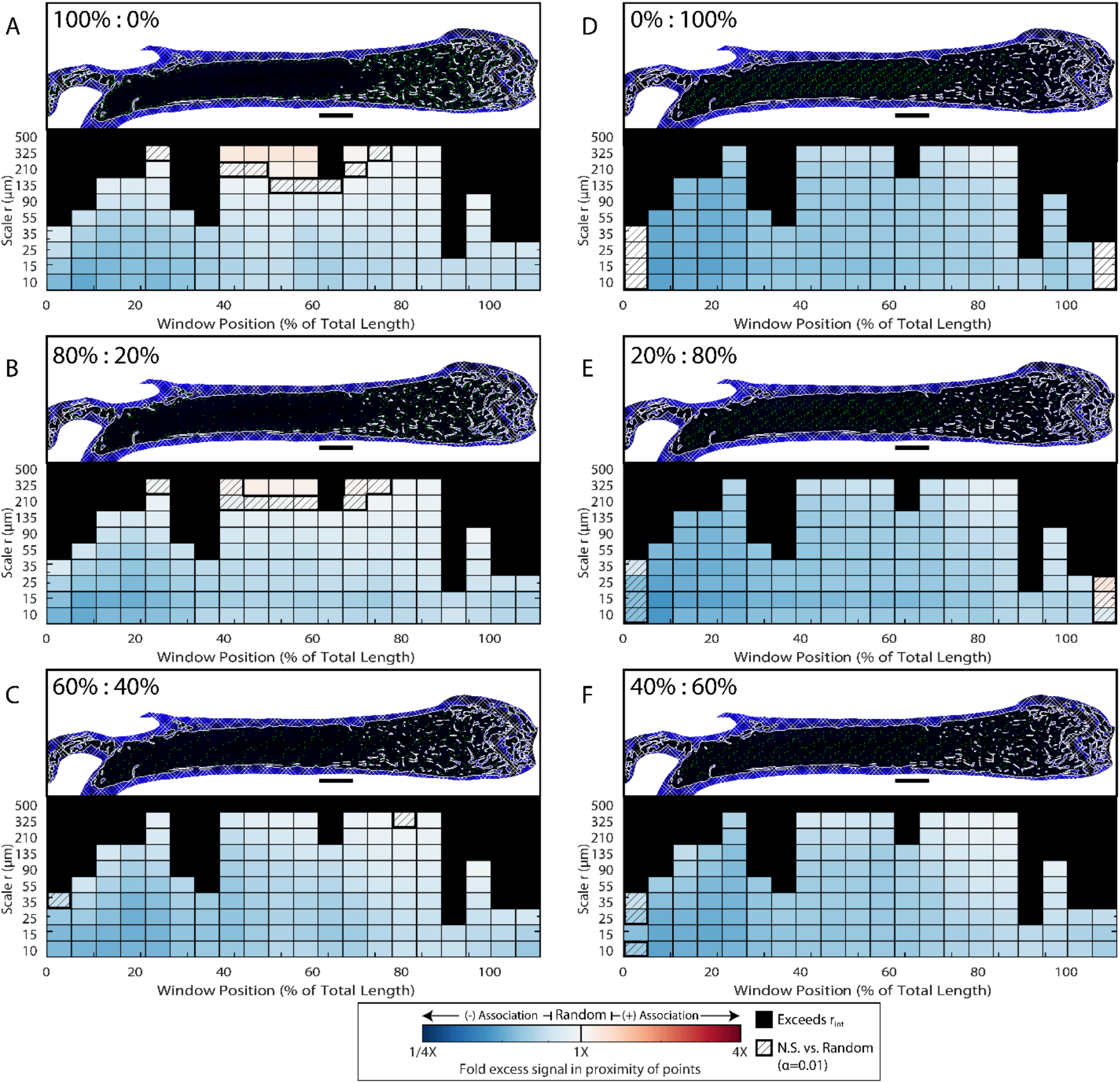
RSRK Testing in Bone. In the same manner of Figure 1B, two regions were defined in which the density of a random point process (green) was manually specified prior to computing RSRK of the full point process with respect to the signal (blue). Region A was defined as within 90 μm of the edge of the endosteum (defined by a binary mask). Region B was defined as any area within the marrow but not within the first region. The test images used for each test are provided. Solid and cross-hashed white areas in the test images indicate the regions in which random points were not placed. The densities of the points in region A vs. region B for each test image are as follows: 100% to 0% (A), 80% to 20% (B), 60% to 40% (C), 0% to 100% (D), 20% to 80% (E), and 40% to 60% (F). 500 μm scale bars are provided in each image. A sliding Ripley’s K plot is shown for each test image. The hue indicates the type of pattern (red=associated, white=random, blue=disassociated), and saturation indicates the strength of the pattern (dark=strong, light=weak). Cross-hatched tiles are not significant (N.S.) versus random but all other tiles are significant versus random (α=0.01, n=199). Black tiles exceed the interpretable scale *r_int_* for the window. A logarithmic scale is used for *r*.

Overall the RSRK results for every test image indicated a significant negative association at small scales, reflecting the fact that the point process and signal never colocalize. The key difference in RSRK results between these test images are visible at scales greater than 90μm. In patterns with point processes that are more dense near the bone (Figure 3 A-C), the RSRK results transition from indicating negative association between the point process and signal at small scales to indicating positive association between the point process and signal, with the shift being most pronounced where point density is 100% in region A and 0% in region B (Figure 3A) and less pronounced where point density is 60% in region A and 40% in region B (Figure 3C). In patterns with point processes that are less dense near the bone (Figure 3 D-F), RSRK detects significant negative association between the point process and signal at scales up to 325 μm in most regions of the bone. This effect was most pronounced in Figure 3 D (0% in region A to 100% in region B) and less pronounced in less extreme cases (Figure 3F).

The ability of RSRK to detect different types of patterns within the context of the bone marrow was also tested. To accomplish this, three test images were prepared: One with point process and signal positively associated and arranged regularly; one with point process and signal negatively associated and arranged regularly; and one with point process and signal positively associated and arranged into clusters (Supplementary Figure S3). RSRK testing of these images indicated the correct type of patterning at small scales in all images tested; significant positive association (Supplementary Figure S3 A,C); and significant negative association (Supplementary Figure S3 B) were observed. Overall, RSRK also detected the correct type of secondary patterning in the test images with patterns at scales ranging from 135 to 225μm with a predicted positive association is predicted from 135 to 225μm (Supplementary Figure S3A and B), while additional positive association (clustering) is indicated from 135 to 225μm (Supplementary S4 C). Notably, there is an inflection point near 225μm where the type of patterning detected shifts from positive association to nearly random (Supplementary Figure S3 C), which corresponds to the approximate radius of the clusters in the image. Taken together with the previous results, RSRK retains the properties demonstrated during verification testing even in complex spatial maps like the bone marrow.

### CXCL12+ cells self-organize into diffuse clusters throughout the bone marrow

To search the bone marrow for spatial structures that may serve as landmarks for the HSC niche, the spatial distribution of CXCL12+ regulatory cells was evaluated with respect to themselves as well with respect to three other major physiological features of the bone marrow: the vasculature, the bone, and LepR+ stromal cells. The type and strength of spatial patterning that exists between CXCL12+ cells and these structures were assessed and the scale at which the patterning occurs was characterized. Identifying the scale of the pattern can allow the prediction of the type of signalling that may be used to coordinate a particular pattern. Small-scale patterns are likely the result of autocrine signalling, a cell signalling to itself, or juxtacrine signalling, a cell signalling to an adjacent cell. Mid-scale patterns are likely the result of paracrine signalling in which a cell secretes a chemical signal that diffuses to distant cells up to a distance of 250μm (Francis and Palsson, 1997). Finally, large-scale patterns, greater than 250μm in scale, are likely the result of endocrine signalling, in which a chemical signal diffuses into the bloodstream and is transported to very distant targets. Altogether this analysis allowed the discovery of spatial structures between HSC-regulating CXCL12+ cells and other bone marrow features.

To evaluate the spatial relationship between CXCL12+ cells and CXCL12 signal in the bone marrow, CXCL12+ cell coordinates were obtained via automatic segmentation of immunofluorescent images of CXCL12 expression in mouse femurs (Supplementary Figure S1A, STAR Methods) as the point process. Results from this analysis were pooled from five mice (STAR Methods). To evaluate self patterning of CXCL12+ cells, the CXCL12+ cell point processes were compared to CXCL12 signal from the same images. RSRK analysis detected positive association between CXCL12+ cells and CXCL12 signal at the autocrine scale throughout the bone marrow (Figure 4B). It was hypothesized that the small scale association between CXCL12+ cells and CXCL12 signal would result in a univariate analysis due to previous validation testing of similar phenomena (Figure 1C). However, RSRK also detected patterning at larger scales. At the juxtacrine scale, RSRK detected up to 1.5 times more CXCL12 signal than expected for a random distribution of cells (Figure 4B) and at the paracrine scale RSRK detected up to 125 percent excess CXCL12 signal in the proximity of CXCL12+ cells (Figure 4B).

**Figure 4.**
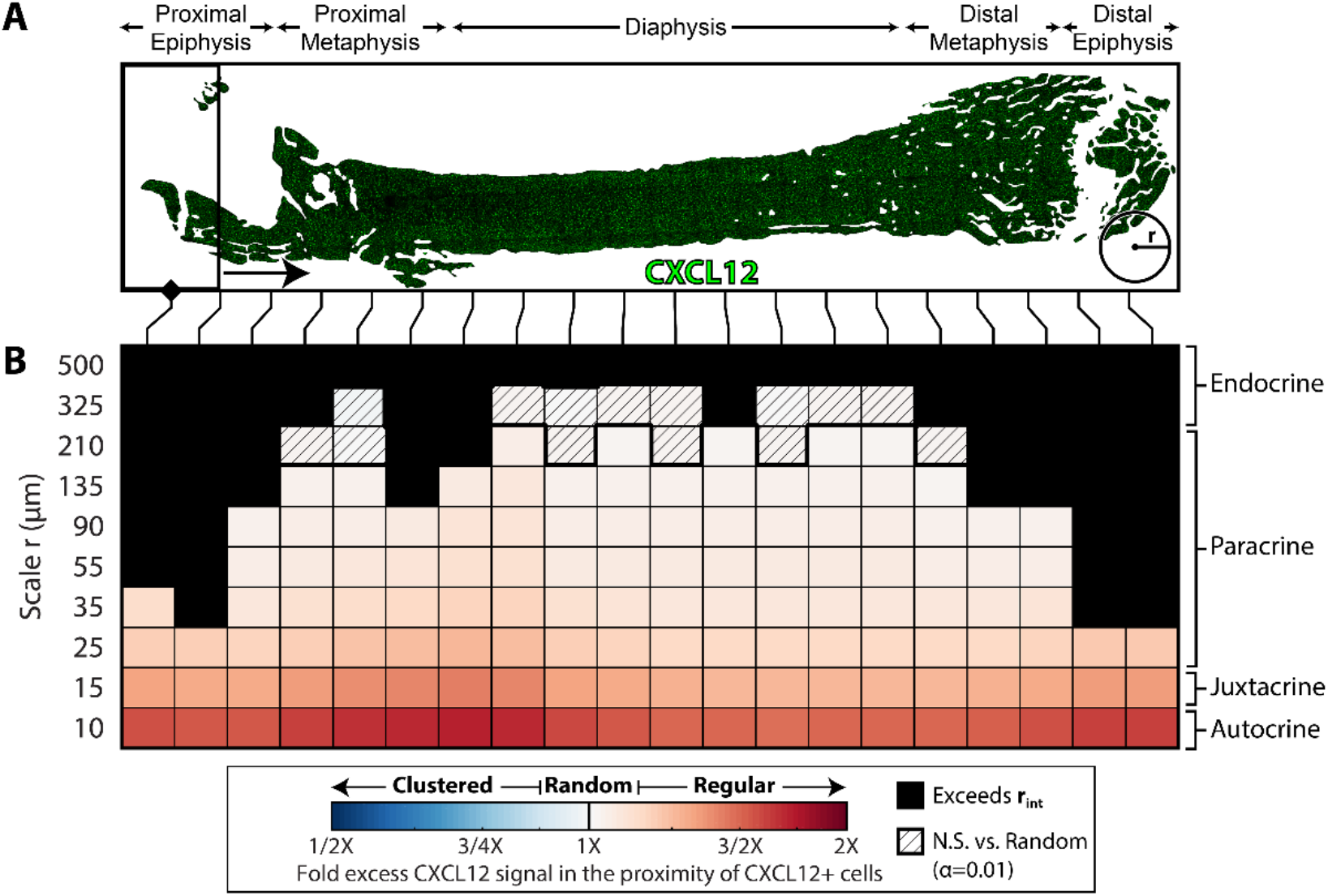
CXCL12+ Cells Self-organize into Diffuse Clusters throughout the Bone Marrow. (A) Masked immunofluorescent image input and rolling window scheme for RSRK analysis of CXCL12+ cells and CXCL12 signal. The window slides from left to right and is shown to scale *n_w_* = 20, *v* = 0.5. Data were pooled from images of five 12-week-old male mouse femurs. The masked area is shown in white. A circle with a 500 μm radius circle is provided in each image for scale. (B) Sliding Ripley’s K plot to evaluate the spatial relationship between CXCL12+ cells and CXCL12 signal. The hue indicates the type of pattern, and saturation indicates the strength of the pattern. Cross-hatched tiles are not significant (N.S.) versus random but all other tiles are significant versus random (α=0.01, n=199). Black tiles exceed the interpretable scale *r_int_* for the window. A logarithmic scale is used for *r*. Estimates for the signaling types associated with a given scale *r* are provided on the right.

It is noteworthy that no distinct regional differences in the type of univariate patterning was detected between CXCL12+ cells. Across the bone, strong clustering between CXCL12+ cells at small scales transitioned to weaker clustering at large scales (Figure 4B). Furthermore, RSRK did not detect any significant disassociation (indicated by Blue in the RSRK results) between CXCL12+ cells and CXCL12 signal (Figure 4B). In summary, RSRK suggests that CXCL12+ cells form clusters throughout the bone marrow at scales possibly indicative of autocrine, juxtacrine, and/or paracrine signaling, and exhibit no spatial regularity at any of the scales measured, implying that CXCL12+ cells self-organize into diffuse clusters throughout the bone marrow.

### The bone has minimal influence on CXCL12+ cell spatial localization throughout the marrow

Bone has long been suspected to be involved in the regulation of HSCs. Compared to perivascular cells, bone cells express 1000 fold less CXCL12 (Morrison and Scadden, 2014). However, bone may indirectly regulate HSCs by coordinating the position of nearby CXCL12+ cells. To test this, RSRK was used to analyze the spatial organization of CXCL12+ cells with respect to Collagen 1 (Col1), which marks bone and the endosteal region. Using multiplexed images of Collagen 1 and CXCL12 expression in the bone marrow, a point process for CXCL12+ cells was segmented and RSRK was used to compare the distribution of this point process to Col1 signal in the bone marrow (Figure 5A). During the simulation of randomness for this particular experiment, random points were permitted only within the bone marrow stroma to reflect the fact that CXCL12+ cells are generally not found within the bone itself.

**Figure 5.**
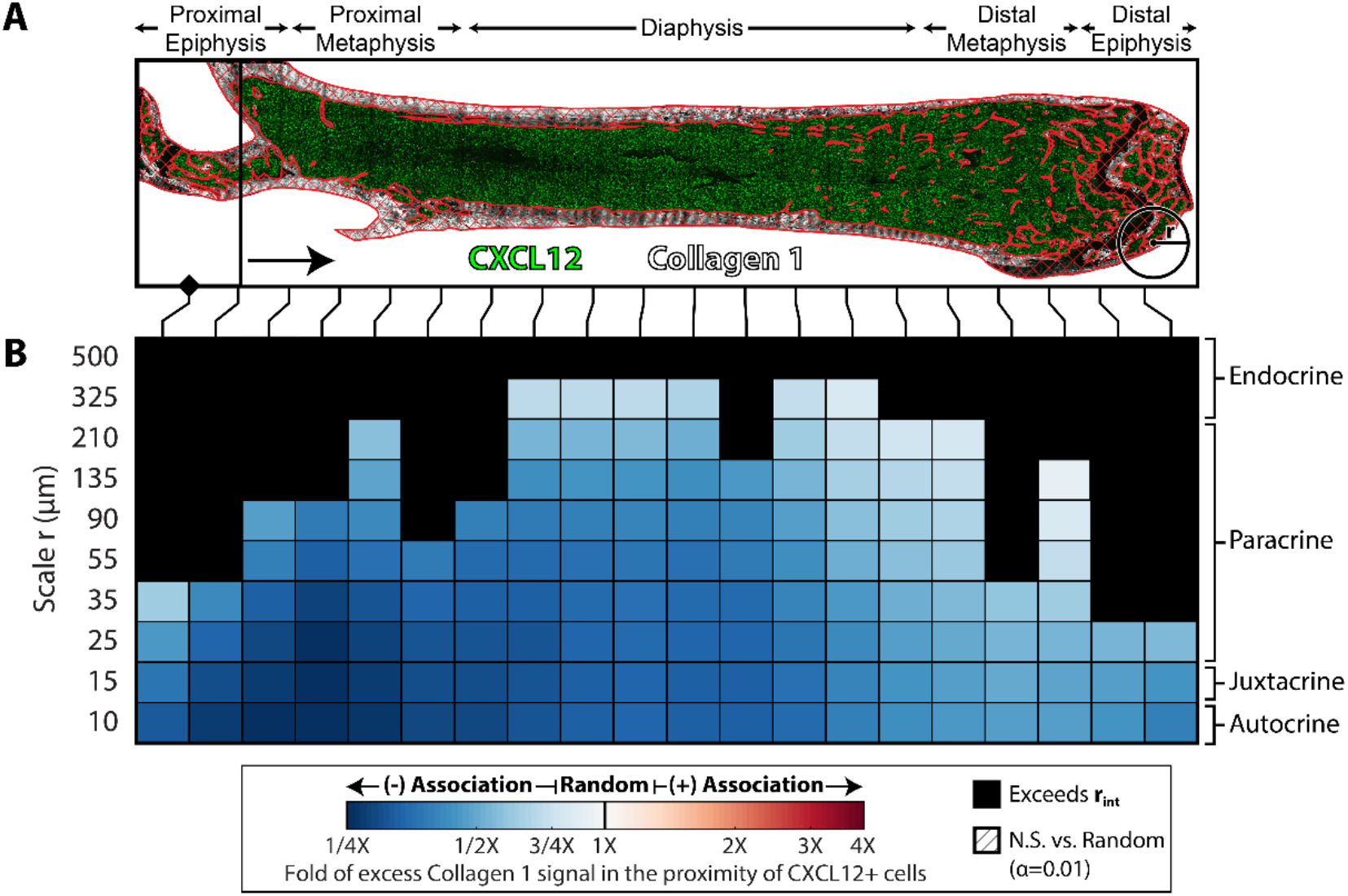
RSRK detects negative spatial association between CXCL12+ cells and bone. (A) Masked immunofluorescent image input and rolling window scheme for RSRK analysis of CXCL12+ cells and collagen-1 signal. The window slides from left to right and is shown to scale *n_w_* = 20, *v* = 0.5. Data were pooled from images of two 12-week-old male mouse femurs. The masked area is shown in white. During the simulation of randomness, random CXCL12+ cells could not fall within the red cross-hashed area. A circle with a 500 μm radius circle is provided for scale. (B) Sliding Ripley’s K plot to evaluate the spatial relationship between CXCL12+ cells and collagen-1 signal. The hue indicates the type of pattern, and saturation indicates the strength of the pattern. Cross-hatched tiles are not significant (N.S.) versus random, but all other tiles are significant versus random (α=0.01, n=199). Black tiles exceed the interpretable scale *r_int_* for the window. A logarithmic scale is used for *r*. Estimates for the signaling types associated with a given scale *r* are provided on the right.

Overall, RSRK detected a significant negative spatial association between CXCL12+ cells and bone. At the autocrine scale, the excess ColI signal in the proximity of CXCL12+ cells was one-quarter of the expected collagen-1 signal under random distribution of CXCL12+ cells (Figure 5B). This negative association at small scales was reminiscent of the strong dissociation noted in the sensitivity tests for RSRK in the bone (Figure 3) and reflects the fact that CXCL12+ cells do not colocalize with bone. Furthermore, negative association between CXCl12+ cells and bone at scales up to about 50μm was observed in all regions of the bone marrow, and at scales up to 500μm in all regions excluding the epiphyses (Figure 5B). Broadly, the RSRK results indicate significant negative association between CXCL12+ cells and bone at small scales that weakens at larger scales suggesting that the bone has little impact on CXCL12+ cell spatial localization in the bone marrow.

### CXCL12+ cells are weakly associated with the perivascular and stromal regions

There is increasing evidence that the vasculature plays a significant role in regulating hematopoiesis, HSCs, and their progeny (Asada et al., 2017; Ding and Morrison, 2013; Duarte et al., 2018; Kunisaki et al., 2013). There is also evidence that LepR+ cells, including LepR+ vasculature, play a role in hematopoietic regulation (Comazzetto et al., 2019; Zhou et al., 2014). To evaluate whether CXCL12+ exhibit spatial association with these important features of the bone marrow, RSRK was used to compare CXCL12+ cell coordinates to CD31 signal, which labels all vasculature in the bone marrow (Coutu et al., 2017), and LepR signal. RSRK analysis indicated that at least a subpopulation of CXCL12+ cells co-express CD31 and LepR or equivalently that CXCL12+ cells and CD31+ and LepR+ cells reside near eachother, as indicated by the assocation between CXCL12+ cells and the two cues at the autocrine scale (Supplementary Figure S4C-D). Throughout the bone marrow, at larger scales, the patterning of CXCL12+ cells with respect to CD31+ vasculature weakened to near randomness starting at scales near 50μm. In the proximal metaphysis only, CXCL12+ cells appear to transition from positively associated with LepR+ cells at scales less than 30μm, to randomly associated, to negatively associated with LepR+ cells at scales greater than 50μm. In all other regions of the bone marrow, CXCL12+ cells appear to be weakly associated with LepR+ cells at all large scales.

## DISCUSSION

The study of morphogenesis, embryogenesis, and spatial pattern formation has been hampered by a lack of robust computational tools to quantify and detect spatial patterning of multiple tissue features. No computational tools exist that are *feature agnostic*, such that spatial patterning can be analyzed between features of any kind regardless of whether they are point, line, or area features. Traditional object-based spatial analyses, like Ripley’s K, can be confounded by inhomogeneous point patterns and non-point like features. By combining object-based and signal-based analyses of spatial organization, we developed Rolling Signal-Based Ripley’s K (RSRK). RSRK permits region-specific analysis of inhomogeneous spatial patterns by implementing a rolling window scheme and incorporates the quantification of signal distribution (as opposed to point pattern distribution) to analyze spatial patterning with respect to non-point like features. By including random simulation in the calculation of RSRK, each result is normalized and ascribed a significance level to allow RSRK to be used to determine the type and scale of a given spatial pattern. Furthermore, RSRK can readily pool results from multiple samples.

To verify the properties of RSRK, it was applied to a large set of pre-defined test images. Using these test images, RSRK could identify different types of spatial patterns and was responsive to subtle changes in spatial patterning. Additionally, it was confirmed that RSRK can be used in univariate analyses and that self-expression does not confound the detection of large-scale patterns. Furthermore, the ability of RSRK to detect regional shifts in spatial patterning and by masking these test images, to remove data, was evaluated and it was confirmed that RSRK was robust to small changes in the geometry of the region being analyzed. A known-characteristic of traditional Ripley’s K statistics is that aspects of the geometry of spatial patterns (e.g. cluster radius, regular spacing) manifest in the Ripley’s K curve as peaks, valleys, and inflection points (Amgad et al., 2015). The verification tests also demonstrated that RSRK shares this important feature of traditional Ripley’s K statistics such that RSRK can be used not only to detect the presence of different kinds of spatial patterns but also to infer various metrics associated with those spatial patterns.

To better understand the interactions within the bone marrow that may drive HSC cell fate decisions and to identify landmarks for the HSC niche, RSRK was used to analyze immunofluorescent images of the mouse bone marrow to characterize the spatial patterning of CXCL12+ regulatory cells with respect to themselves as well as to CD31+ vasculature, Collagen 1+ bone, and LepR+ stromal cells. The spatial localization of known HSC regulatory cells, specifically CXCL12+ cells, can further elucidate where HSCs are more likely to localize and identify mechanisms that may influence that localization.

By performing univariate RSRK between CXCL12+ cells and CXCL12 signal, it was discovered that CXCL12+ cells display positive associations with the CXCL12 signal that are attractive versus random distribution (Figure 4). At small scales this strong positive association is due to self expression. To rule out that this alters or masks larger scale patterning, verification testing of similar phenomena were done and demonstrated that self-expression does not mask larger scale patterning (Figure 1C). At scales greater than 20μm, RSRK predicts moderate spatial association that nears randomness as scale increases (Figure 4). This is similar to results from the test images of self-expression results (Figure 1C), in which points that express signal are arranged in clusters at large scales, and RSRK indicated strong association that weakened as scale increased. Altogether, these results suggest that CXCL12+ cells form clusters with other CXCL12+ cells throughout the bone marrow. These, are the first spatial structure identified within the bone marrow that express critical HSC regulating cues, making them a candidate landmark for the elusive HSC niche.

Early studies of the bone marrow suggest that primitive HSCs reside near the endosteum and that their progeny reside closer to the center of the marrow cavity and that perturbations of the osteoblasts, found in the endosteum, could influence HSC density in the bone marrow (Sugiyama et al., 2018). However, later studies found that HSCs preferentially localized near sinusoidal vasculature instead of the endosteum (Acar et al., 2015; Kiel et al., 2005). Acar *et al*. also determined that the distribution of HSCs is significantly reduced near the endosteum in the diaphysis (Acar et al., 2015). This same study also determined that HSCs are significantly rarer in the epiphysis and metaphysis where the trabecular bone is most dense versus the diaphysis where there is very little trabecular bone (Acar et al., 2015). Corresponding patterning among CXCL12+ cells with respect to bone could suggest that CXCL12+ cells are involved in the spatial patterning that occurs between HSCs and bone. RSRK analysis indicated a negative association between CXCL12+ cells and collagen 1 signal (bone) that was exceptionally significant versus random. 50% to 75% less collagen 1 signal was detected within 10μm of CXCL12+ cells versus when CXCL12+ cells were placed randomly within the marrow cavity (Figure 5). The strength of this negative association weakened at larger scales suggesting far less signal observed in the proximity of CLCL12+ cells than was expected from random simulation. At first glance, this would suggest that CXCL12+ cells preferentially localize away from bone, however, the verification results from RSRK testing in bone temper these findings (Figure S3) because point processes with different densities were manually defined in two different regions, one near the bone and one far from the bone, and then RSRK was computed (Figure 3). In terms of the scale at which a given type of spatial patterning is detected, the results for RSRK analysis of CXCl12+ cells with respect to bone are most similar to the verification tests in which the point process density was reduced near the bone (Figures 3 D-F). RSRK applied to all 3 of these images, predicted negative spatial association between point process and signal at all interpretable scales that approached random as scale increased.

Perivascular endothelial cells express CXCL12 and are necessary for HSC maintenance (Comazzetto et al., 2019; Ding and Morrison, 2013; Ding et al., 2012; Greenbaum et al., 2013). CD31 labels all vascular elements of the bone marrow and strongly labels sinusoids (Coutu et al., 2017). Previous analyses of the spatial correlation of CXCL12 and CD31 vascular signals considered only whether two factors were coexpressed. These studies found that many perivascular and endothelial cells express large amounts of CXCL12 (Coutu et al., 2017; Ding and Morrison, 2013). However, when bivariate RSRK was used to evaluate the patterning of CXCL12+ cells with respect to CD31+ vasculature, the spatial association was barely significant versus random and only consistent at scales less than 25μm throughout the bone marrow. At most, 25% excess CD31 signal was detected in the proximity of CXCL12 cells (Supplementary Figure S4A,C). These results do not disprove previous studies but suggest that vasculature may be so replete in the bone marrow that any cell, including CXCL12+ cells, if randomly placed in the bone marrow, may appear to coexpress large amounts of CD31. These results also suggest that CXCL12+ cells preferentially localize near CD31+ cells but also that this preferential localization is very subtle and only impactful at small scales.

Like vasculature, LepR+ cells have been reported to play a significant role in HSC regulation (Asada et al., 2017; Comazzetto et al., 2019; Ding and Morrison, 2013) and to co-express CXCL12 (Coutu et al., 2017; Ding and Morrison, 2013). Coutu et al. report 45% of CXCL12+ cells also express high levels of LepR (Coutu et al., 2017). The positive association between CXCL12+ cells and LepR signal at small scales corroborates these findings (Supplementary Figure S4 B and D). However, the RSRK results for CXCL12+ cells with respect to LepR+ cells indicated a relatively weak positive association when compared to random at most spatial scales and regions of the bone suggesting that CXCL12+ cells are loosely attracted to LepR+ cells. The only exception was in the proximal metaphysis where negative association between CXCL12+ and LepR signal was weak versus random, which was detected at scales greater than 35 μm. Like with CD31 + vasculature, these spatial associations that are weak versus random suggest that the placement of LepR+ cells may only have a subtle influence on CXCL12+ cell organization.

Hematopoietic stem cells (HSCs) are believed to be maintained and regulated through the concerted effort of a variety of cells and regulatory cues collectively called the HSC niche. Our data suggests that HSC niches appear to be limited and that HSCs must compete for niche sites to be maintained (Post and Clevers, 2019; Schofield, 1978; Shimoto et al., 2017). Until now, the bone marrow research community has not identified a structure that demonstrates the HSC niche (Wei and Frenette, 2018). In this study, RSRK was used to identify CXCL12+ cell clusters that we hypothesize to be niche sites for which HSCs compete. Because CXCL12 is the only known chemoattractant for HSCs (Wright et al., 2002), clusters of CXCL12 expressing cells will create a gradient of CXCL12 that are hypothesized to draw CXCL12-sensitive HSCs toward the center of the CXCL12+ cell cluster (Figure 6). Therefore, the center of the cluster would have the highest concentration of maintenance and retention cues expressed by CXCL12+ cells representing an ideal site for HSC maintenance retention. Differentiation and proliferation of HSCs can also be explained under this hypothesis. Since HSCs detect and respond to gradients in CXCL12 via the CXCR4 receptor (Wright et al., 2002). Studies have shown that the loss of CXCR4, and therefore loss of CXCL12+ sensitivity, occurs during HSC differentiation (Sugiyama et al., 2006), thus causing a differentiating HSC to disassociate from the niche composed of CXCL12+ cells, and eventually result in their differentiation (Figure 6).

**Figure 6.**
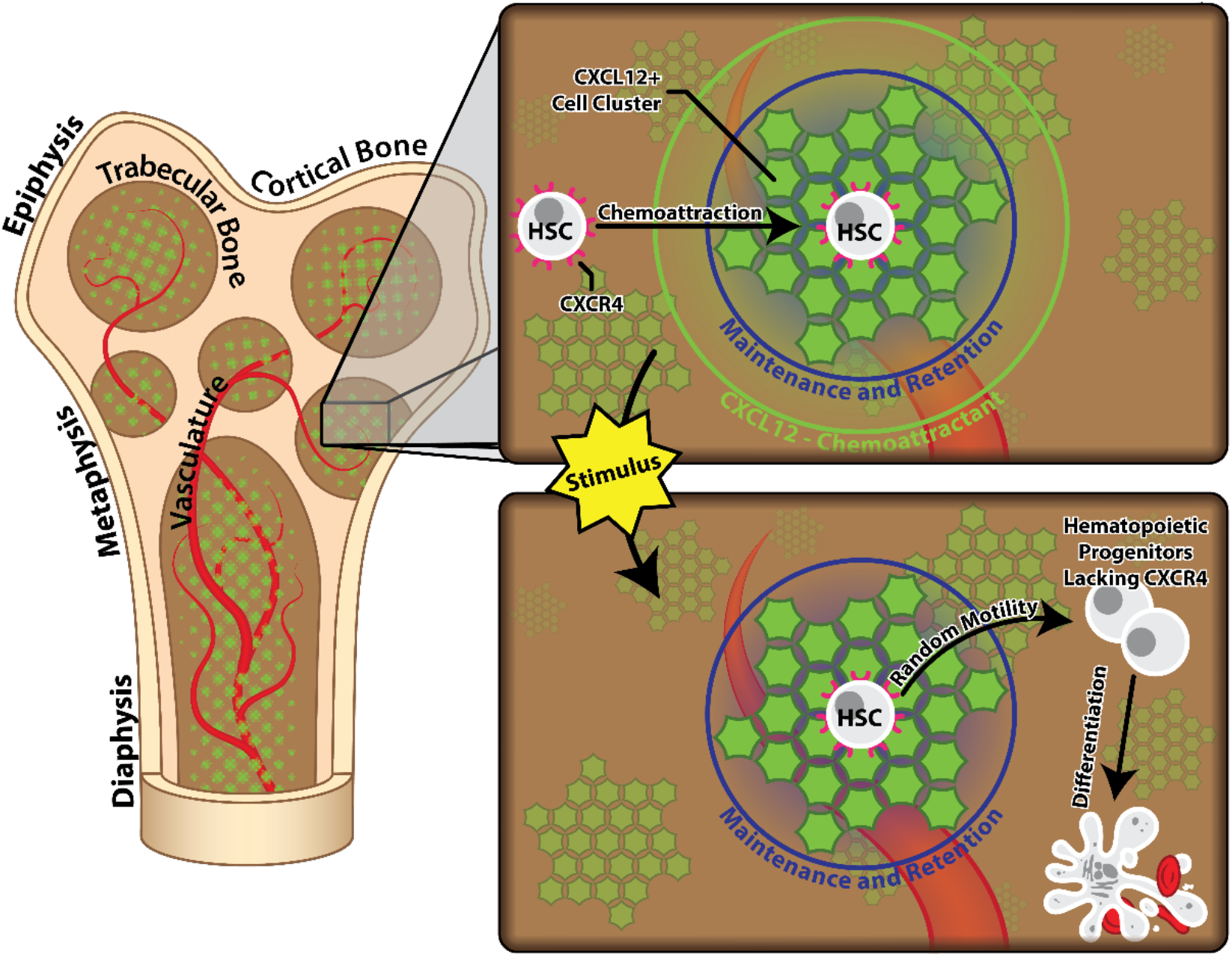
A Hypothesis: HSC regulation is achieved through control of CXCL12 expression and sensitivity. The presence of CXCL12+ cell clusters in the bone marrow suggest that HSCs are regulated extrinsically through their exposure to CXCL12 chemoattractant and intrinsically by their sensitivity to CXCL12 chemoattractant. Clusters of CXCL12+ cells in the bone marrow may create gradients in CXCL12 that attract CXCR4-expressing HSCs where they are exposed to the highest concentration of maintenance and retention cues. An external stimulus that disrupts CXCL12 expression by CXCL12+ cells or CXCR4 expression by HSCs would cause HSCs to disassociate from the niche and lose exposure to maintenance and retention cues, eventually leading to their proliferation and differentiation.

In conclusion, we have developed a robust computational tool, RSRK, for spatial analysis of histological specimens. RSRK combines object-centric and signal-centric analyses of spatial correlation with a moving window scan to characterize the spatial structure between point and non-point like features across an inhomogeneous region. In this study, RSRK, was used to identify previously undescribed landmarks and patterns of cell distribution in the bone marrow. In particular CXCL12+ cell clusters were identified within the bone marrow, enabling a hypothesis for how these cell clusters could form the basis for the HSC niche. This identification of cell patterns in the bone marrow provide insights for the basis of the elusive HSC niche highlighting the utility of RSRK as a tool of both analysis and discovery that can broadly be applied to the study of tissue and organism morphogenesis.

## METHODS

### Key Resources Table

| RESOURCE | SOURCE | IDENTIFIER |
| --- | --- | --- |
| **Software and Algorithms** |  |  |
| MATLAB version 9.5 | The MathWorks, Inc. 2018 | https://www.mathworks.com |
| MATLAB Image Processing Toolbox version 10.3 | The MathWorks, Inc. 2018 | https://www.mathworks.com/products/image.html |
| FIJI (ImageJ 1.52p) | (Schindelin et al., 2012)<br>(Schneider et al., 2012) | https://fiji.sc/ |
| RSRK Toolbox | This paper | https://github.com/cphealy8/RSRK |
| **Source Images** |  |  |
| Murine Bone Marrow Images | (Coutu et al., 2017) | http://dx.doi.org/10.1038/nbt.4006 |

## LEAD CONTACT AND MATERIALS AVAILABILITY

Further requests for information and resources should be directed to and will be fulfilled by the Lead Contact, Tara Deans

## METHOD DETAILS

### Image Acquisition

Immunofluorescent Z-stacks were generously provided by Dr. Timm Shroeder (Coutu et al., 2017). The images we used in our analysis corresponded to Figures 3C, 5C-E, and 6D from their 2017 paper, *Three-Dimensional Map of Nonhematopoietic Bone and Bone-Marrow Cells and Molecules* (Coutu et al., 2017). Pixels in these images had widths and heights equal to 0.69μm.

### Projection and Masking

Raw z-stacks from the Coutu database were first converted (at maximum resolution) to 2-dimensional images using ImageJ’s built-in maximum intensity projection utility and using the full width of the stack. Binary masks to distinguish bone marrow from the bone in each image were generated manually using Photoshop. These masks were used to define the region analyzed via RSRK, and the region in which points are randomly placed during random simulation. For bone marrow analysis, the region to be analyzed and in which random simulation occurs was specifically the bone marrow stroma. This excluded regions corresponding to cortical and trabecular bone and regions in the image that were outside the bone. Importantly, for the analysis of CXCL12+ cell distribution with respect to Collagen 1+ bone, two masks were defined. The first defined the region in which analysis would occur and included the bone marrow stroma as well as bony regions such as the cortical and trabecular bone. The second mask defined where random simulation of CXCL12+ cell position would occur and only included the bone marrow stroma. Full-color images were saved as portable network graphics (PNG) at their full resolution. Masks were saved as bitmaps (BMP) at full resolution. Both immunofluorescent images and masks were downsampled to 10% of there original resolution to a scale of 6.9μm per pixel before being analyzed with RSRK to save on computational time.

### Segmentation and Feature Identification

The point processes used in the density, nearest neighbor, and Ripley’s K analyses were generated via semiautomatic segmentation implemented using binary and morphological operations and a custom MATLAB GUI. First, each image is converted to a single-color channel image to isolate a specific feature (e.g., green for CXCL12). Next, the image is thresholded to isolate a desired range of pixel intensities by specifying both a minimum and maximum pixel intensity. Following thresholding, morphological operations are applied to refine the selected pixels. The morphological operations utilize a disc-shaped kernel with a user-specified radius. Two additional operations are included in our GUI: morphological opening (which omits isolated pixels) and morphological closing (which fills holes in pixel objects). The order in which the operations are applied does matter. The GUI is interactive and permits the user to refine their selection to select and segment specific features in the image (e.g., CXCL12+ cells). Next, the selected features are converted into a binary mask and the individual objects in each mask were analyzed to obtain the coordinates of their centroids and their area. Additional features can be omitted from the set of cell coordinates based on their size using the area controls. Following segmentation, the coordinates of each feature in the image were saved. Finally, the saved feature coordinates were combined with the binary masks from the previous step to define a point process (a set of points and the area in which they occur). The point process is used in the calculation of the bone proximity, nearest neighbor, and Ripley’s K statistics.

To obtain the point processes of CXCL12+ cells used in this study, images from the Coutu database depicting CXCL12 expression were segmented using mySegment.m. A raw image was first flattened, and downsampled, as described previously. First, mySegment.m was used to isolate the channel corresponding to CXCL12 from the image. Second, the thresholding feature of mySegment.m was used to select pixels with intensities in the range of 150 to 255. Third, the morphological operation feature of mySegment.m was used to first apply a morphological close with a kernel radius of 2, followed by a morphological open with a kernel radius of 2, to better refine the selection and remove small pixel noise. No area operations were applied. Finally, mySegment.m converted the masked CXCL12+ cells into a point process. Although mySegment.m does permit manual alteration of the point process, the point processes used in this study were output to be used in RSRK without any manual alterations.

### Data Pooling

The RSRK data from several mice were pooled to produce combined RSRK results shown in this paper. For CXCL12 to CXCL12 RSRK analysis, data were pooled from 5 mice, corresponding to point processes obtained from the CXCL12 channel from images 3C, 5C, 5D, 5E, and 6D of the Coutu database. For CXCL12 to CD31 RSRK analysis, RSRK data were pooled from two mice corresponding to images 3C and 5D of the Coutu Database. For CXCL12 to Collagen 1 RSRK analysis, data were pooled from two mice corresponding to images 5E and 6D of the Coutu database.

To pool RSRK data let there be *M* separate RSRK statistics, the pooled RSRK count *E_M_* is given by

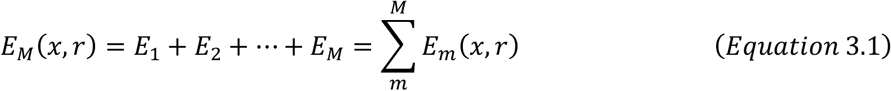

The form of the pooled density *λ_M_* depends on the type of Ripley’s K analysis. For RSRK, *λ_M_* is given by

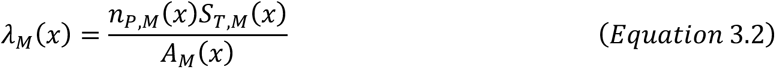

where *n_a,M_*, *S_T,M_*, and *A_M_* are given by

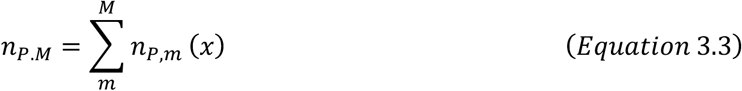

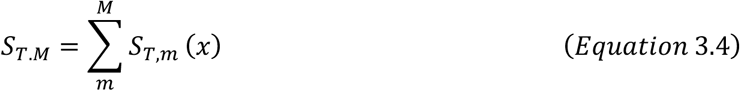

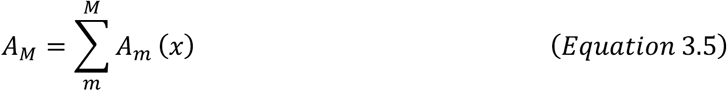

Finally, the pooled RSRK statistic, *K_M_*(*x,r*) is given by

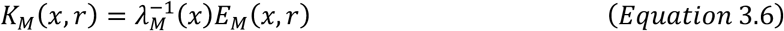

The same approach can be used to pool the results from random simulation to compute a pooled *K_RND,M_*. To pool RSRK statistics for the bone marrow, the window positions *x* analyzed must be the same between samples and must be expressed as a fraction of bone length. The scales *r* that are analyzed and the window parameters *n_W_* and *ν* must also be the same for all samples.

We developed a MATLAB function (**PoolRK.m**) to pool RSRK samples automatically see (https://github.com/cphealy8/RSRK).

### Using RSRK Toolset (step by step)

1. Download and unzip the RSRK toolbox. We recommend working within default RSRK file structure to avoid reference errors.
2. If starting with a confocal z-stack, prepare a flattened 2D image using ImageJ’s built-in projection tools. In ImageJ select Image>Stacks>Z-Project. Then specify slice range and projection type (Maximum Intensity). (See Projection and Masking). Save the flattened image in the directory **../data/images.**
3. Prepare binary masks to define the RSRK analysis regions. Open the flattened image from step 1 in Photoshop, create a new layer atop the original image where the mask will be defined. Lock the original image layer. Do not crop or resize the image at anytime to ensure that the mask and flattened image have consistent resolution. Using the selection, paintbrush, or pen tools define the boundary between the masked and unmasked areas. Next fill the areas to be analyzed (unmasked) with white using the paint bucket tool, and the areas to be excluded from the analysis (masked) with black. Merge all layers so that you only have the mask, then change the image mode to grayscale (Image>Mode>Grayscale), then to Bitmap (Image>Mode>Bitmap). Finally save the image as a.bmp file in the directory **../data/images**.
4. Obtain the desired point process by applying **MySegment.m** to the flattened image from step 1. (See Segmentation and Feature Identification). When the GUI terminates it will automatically prompt you to save the point process and segmentation data. Save the point process under the directory **../data/FullPts**.
5. Isolate the signal image that you wish to analyze. This should be a grayscale image showing only the expression of one marker e.g. CXCL12. This can be easily accomplished by running ImageLoad.m, loading a desired full color image, selecting the channel that corresponds to your target signal, then finally saving the resulting image as a grayscale.png file in the directory **../data/images**.
6. Open **MovingRK_Bivariate_Pts2Signal.m**. The first few lines of this script define the parameters of the RSRK analysis including the downsampling percentage, image scale, image units, window overlap *v*, number of windows *n_w_*, and the scale vector *r*. You can change these if you wish.
7. Run **MovingRK_Bivariate_Pts2Signal.m**. The code will automatically prompt you to select a point process file, a signal image, and a mask to use in the RSRK analysis. It will then automatically begin performing the RSRK analysis and will display the progress of the RSRK calculation.
8. Once the calculation is complete, the script will automatically save the results to **../data/MovingRK**.
9. Optional: Pool RSRK results. Repeat steps 1 through 7 until you have computed all of the individual RSRK results that you wish to pool. Run **PoolRK.m**. The function will automatically prompt you to select an RSRK data file it will then ask if you wish to add more RSRK data to be pooled. Keep adding data until all of the data that you wish pool has been selected beforing selecting “No” the next time the code prompts you to add more data. The code will then automatically pool the data that you selected. Finally, the code will prompt you to save the pooled data.
10. To plot and visualize the results of RSRK analysis, run **MovingRKPlot.m**. The code will automatically prompt you to select the RSRK data you wish to visualize. MovingRKPlot.m will display the results and automatically save them as.eps files to **../results/MovingRKPlots**. It will save two files, one that shows the actual RSRK values and one that shows the significance level of the RSRK values. The colors in the latter correspond to the following significance values and pattern types: Blue, significant disassociation α=0.01; Green, significant disassociation α=0.05; Light Yellow, not significant versus random; Orange, significant association α=0.05; Red, significant association α=0.01.
11. To compute *r_int_* for each mask used in your analysis, open **MaskAnalysis.m**, be sure the variables for image scale, window overlap *v*, number of windows *n_w_*, and *r* match those used in your analysis, then hit **run.** The code will automatically prompt you to select a mask image to analyze, then output a vector **rint** that indicates *r_int_* for each window.

## Acknowledgements

The authors gratefully acknowledge the funding sources to support this work from the National Science Foundation CAREER Program (CBET-1554017), the Office of Naval Research Young Investigator Program (N00014-16-1-3012), the National Institutes of Health Trailblazer Award (1R21EB025413-01), the National Institutes of Health Director’s New Innovator Award (1DP2CA250006-01).

## Author contributions

CPH and TLD conceived the idea of using computational approaches to map the bone marrow. CHP, FRA and TLD designed the experiments. CHP and FRA developed the models and performed formal analysis. CHP conducted all of the computational simulations. CPH wrote the paper. FRA, and TLD reviewed and editied the the paper.

## DATA AND CODE AVAILABILITY

The RSRK toolbox softwareis available in the GitHub repository: https://github.com/cphealy8/RSRK The data used in our analysis is available from the authors upon request.

## Supplemenary Text

### Semi-automatic Image Segmentation

In an effort to automate image segmentation and enhance pattern identification from images, a software package was compiled to generate some of the inputs neded to compute RSRK and compute the RSRK statistic. To generate the input point process needed for RSRK analysis, custom semi-automatic image segmentation software was developed. Given a multiplexed image (Supplementary Figure S1A), this software allows a user to first select which feature they wish to convert into a point process by selecting the corresponding channel in the input immunofluorescent image or, if desired, a combination of features (Supplementary Figure S1B). Next, the user can define a range of pixel intensities to generate a binary mask of their selection (Supplementary Figure S1C). Within the software, the user can further refine this selection via morphological opening (erosion followed by dilation of the selection to remove single pixel noise) and closing (dilation followed by erosion of the selection to fill small holes in the selection) (Supplementary Figure S1D). Lastly, the software allows the user to specify limits on specific morphological aspects of the binary objects in their selection, including maximum and minimum area and solidity. Given the user-defined selection, the software then automatically computes the centroids of the selected features (Supplementary Figure S1E). Finally, the software permits the user to manually edit the point process automatically generated in the previous steps, although this feature of the semi-automatic segmentation software was not used in this study. It should be noted that this image software implements RSRK as a moving window scheme, which is inherently directional. Depending on the direction the user defines, RSRK will produce different results. For this reason, users must apply RSRK in the direction along which they expect the most regional heterogeneity. However, it is important to note that users can apply RSRK to spatially homogenous images, provided they omit the moving-window scheme to analyze the entire region as a whole. Full region analysis with RSRK can be accomplished by setting the window parameters *n_w_* and *v* to zero and 1, respectively. Regardless, RSRK has proven to be an effective tool for analysis and discovery of spatial patterns in tissues. Between regions and experimental groups, RSRK can only compare the type and scale at which a particular spatial pattern occurs.

## Supplementary Figures

**Supplementary Figure S1.**
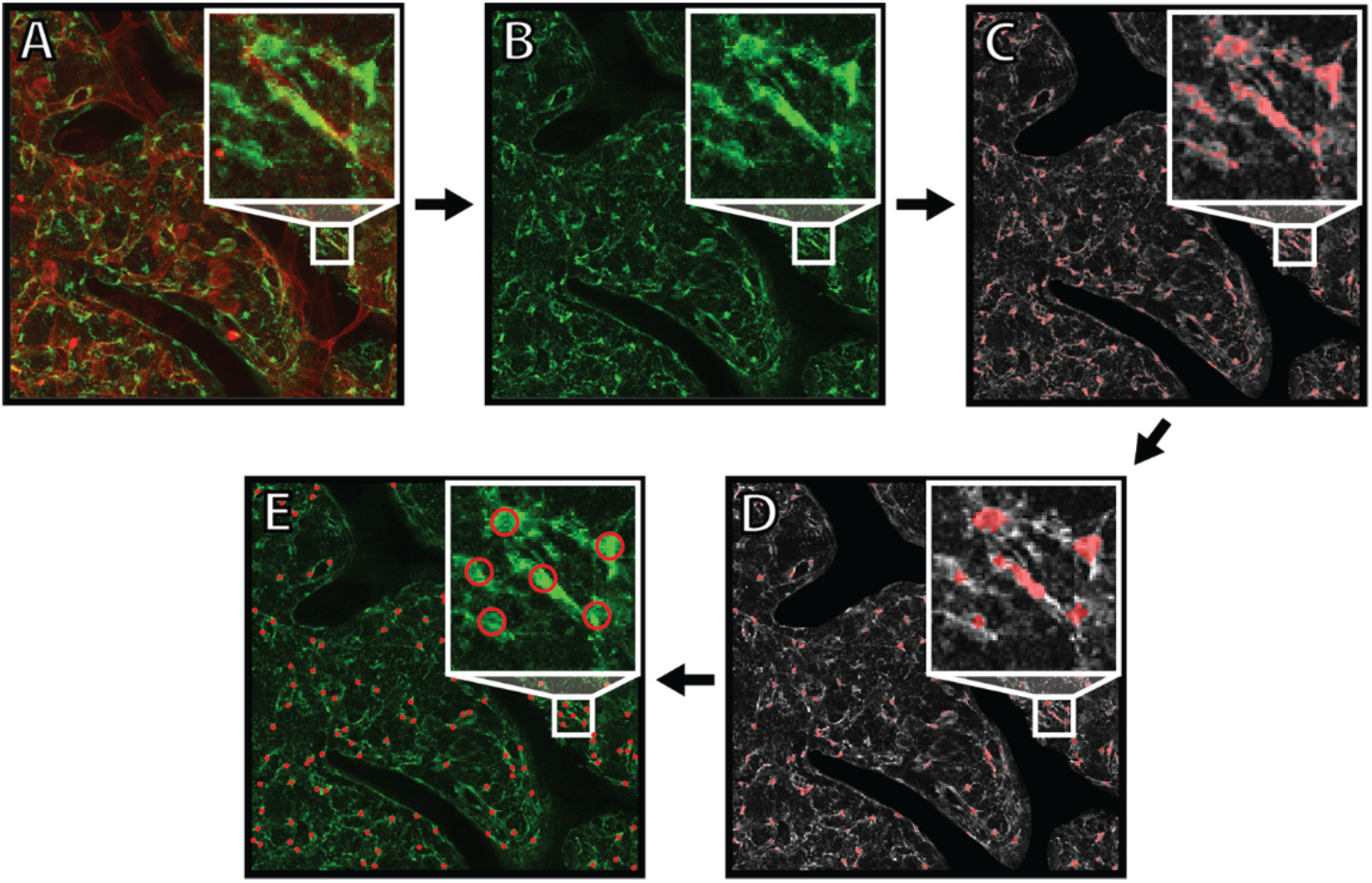
Semi-automatic Segmentation Pipeline. (A) Original Image. CXCL12-GFP Green, CD31 Red. (B) Isolated green channel to segment CXCL12+ cells. (C) Green channel converted to grayscale with binary selection after thresholding in red. (D) Refined selection after morphological operations. (E) Final point process of CXCL12+ cells computed from the centroids of the selections in E.

**Supplementary Figure S2.**
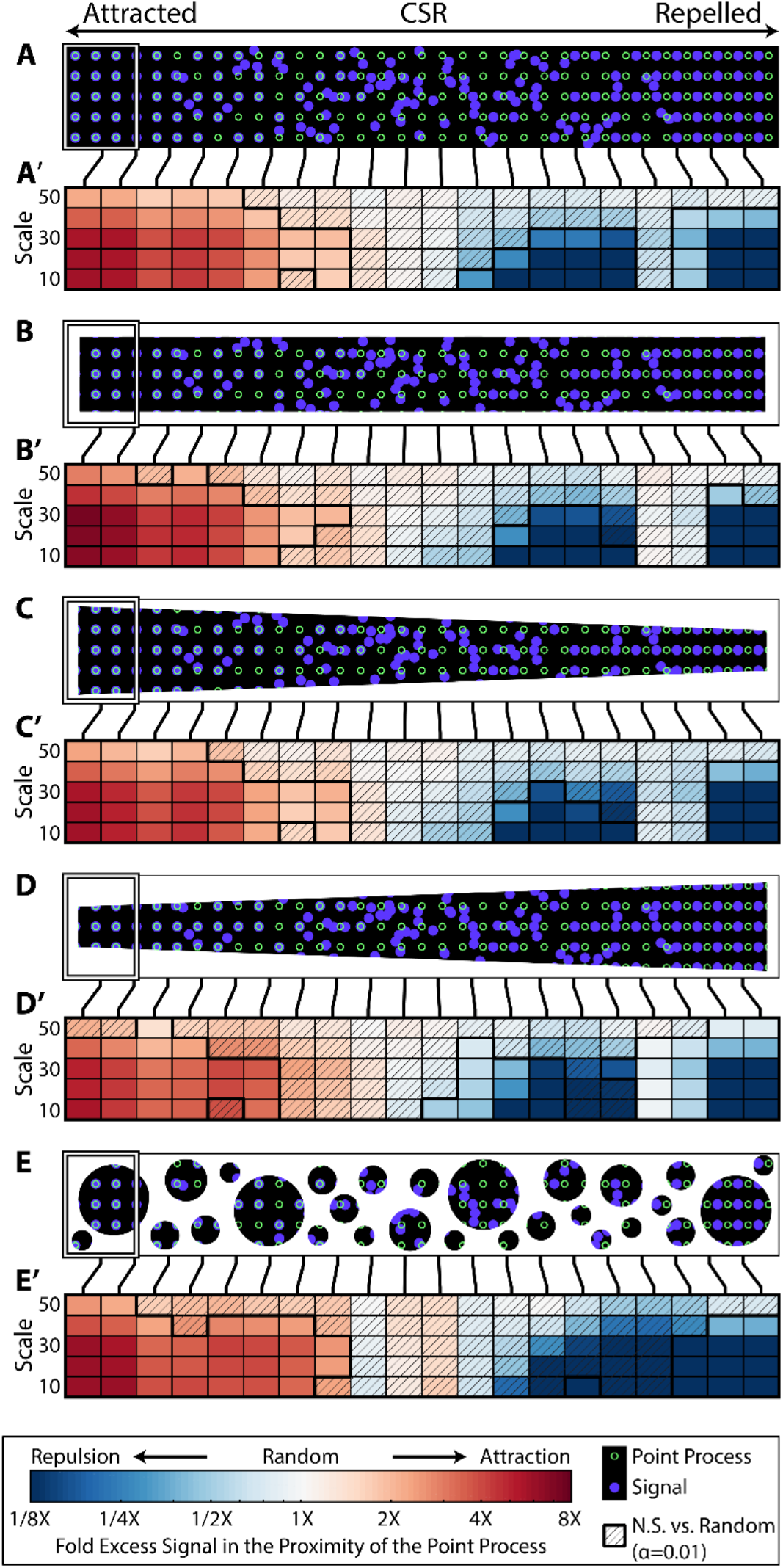
Additional RSRK Masking Validation. (A-E) Additional masked images and point processes used to test RSRK. From left to right, the point process (green) and the signal (blue) transition from perfectly associated, to randomly distributed, to perfectly disassociated. The mask applied to the image is shown in white. No mask was applied in A. The white rectangle shows an example window. (A’-E’) Additional RSRK plots for the images and point processes in (A-E). The hue indicates the type of pattern, and saturation indicates the strength of the pattern. Cross-hatched tiles are not significant (N.S.) versus random. All other tiles are significant versus random. (α=0.01, n=199). Each column corresponds to the RSRK statistic for the window centered at the line connecting the column in (A’-E’) to the images in (A-E).

**Supplementary Figure S3.**
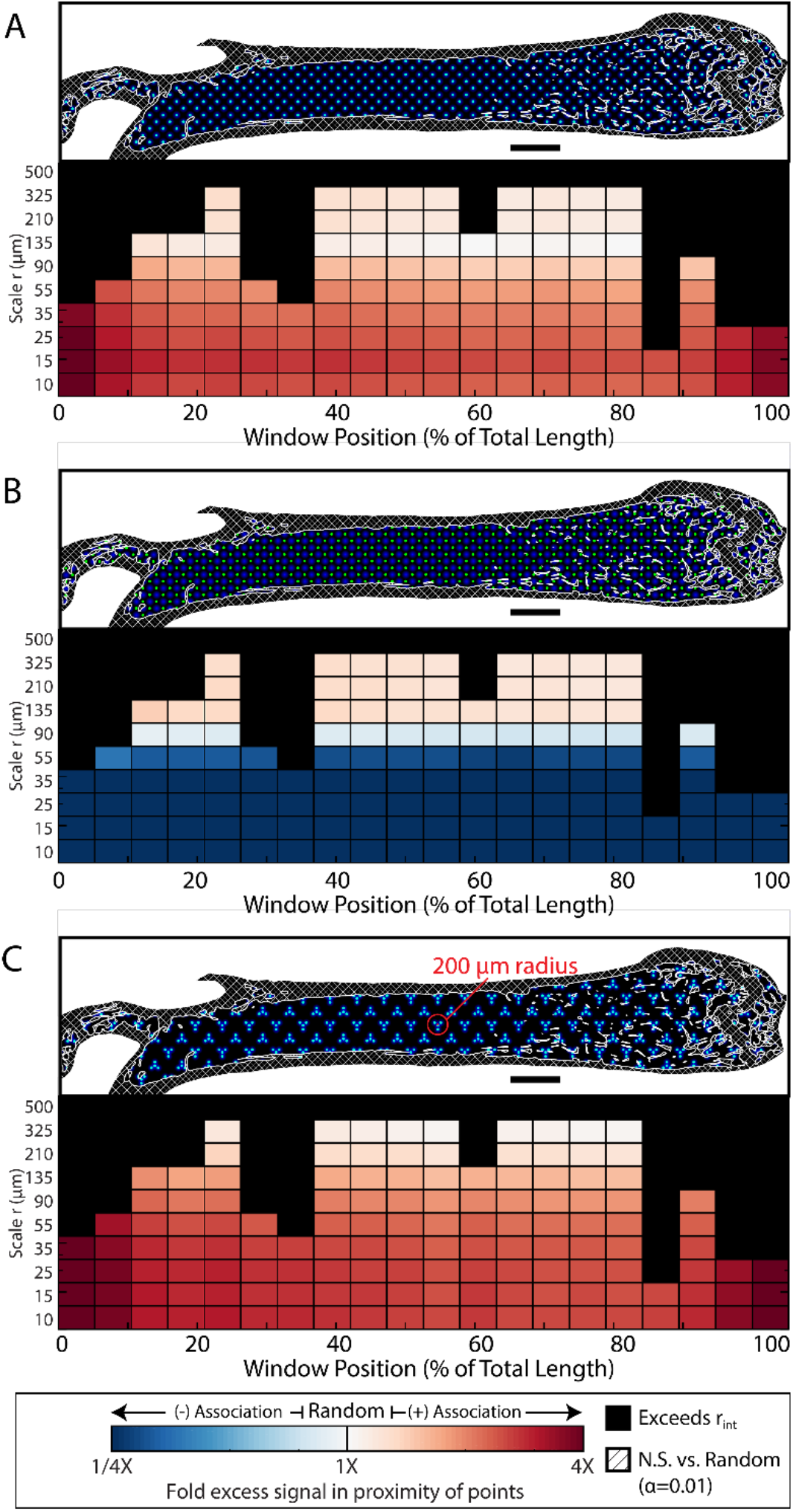
RSRK Specificity Testing in Bone. Pre-defined patterns between a point process and a signal were imposed within the mask for a murine bone then RSRK was computed. The test image used for each analysis is provided and shows the point process (green), signal (blue), and mask used in the calculation of RSRK (Cross-hashed white areas). A sliding Ripley’s K plot is shown for each test image. The hue indicates the type of pattern (red=associated, white=random, blue=disassociated), and saturation indicates the strength of the pattern (dark=strong, light=weak). Cross-hatched tiles are not significant (N.S.) versus random but all other tiles are significant versus random (α=0.05, n=199). Black tiles exceed the interpretable scale *r_int_* for the window. A logarithmic scale is used for *r*. A) Signal and point processes are perfectly associated and otherwise arranged uniformly, B) Signal and point processes are perfectly disassociated and otherwise arranged uniformly, and C) Signal and point process are perfectly associated and arranged in clusters.

**Supplementary Figure S4.**
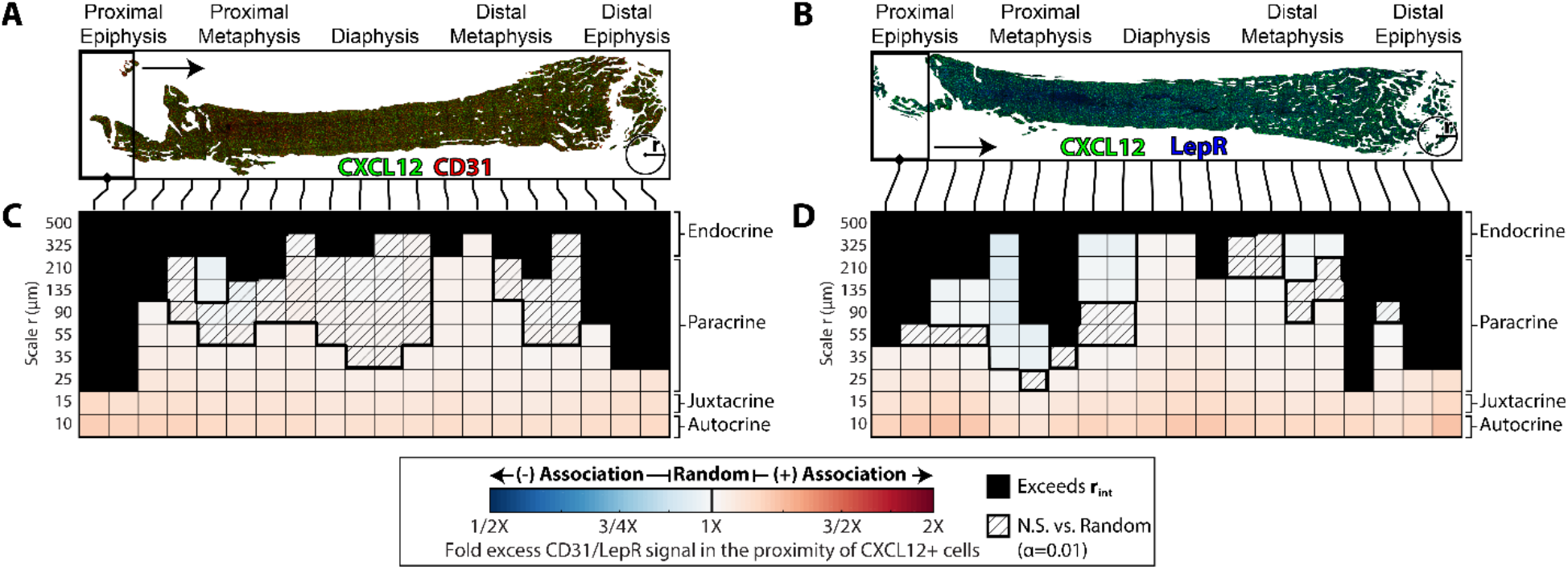
CXCL12+ cells are positively associated with CD31/LepR signal throughout the bone marrow. (A-B) Masked immunofluorescent image input and rolling window scheme for RSRK analysis of CXCL12+ cells and CD31 signal(A) and LepR signal (B). The window slides from left to right and is shown to scale *n_w_* =20, *v* = 0.5. The masked area is shown in white. A circle with a 500 μm radius circle is provided in each image for scale. The data in A were pooled from images of two 12-week-old male mice. (B) Sliding Ripley’s K plot to evaluate the spatial relationship between CXCL12+ cells and CD31 signal (C) and LepR signal (D). The hue indicates the type of pattern, and saturation indicates the strength of the pattern. Cross-hatched tiles are not significant (N.S.) versus random but all other tiles are significant versus random (α=0.01, n=199). Black tiles exceed the interpretable scale *r_int_* for the window. A logarithmic scale is used for *r*. Estimates for the signaling types associated with a given scale *r* are provided on the right.

**Supplementary Figure S5.**
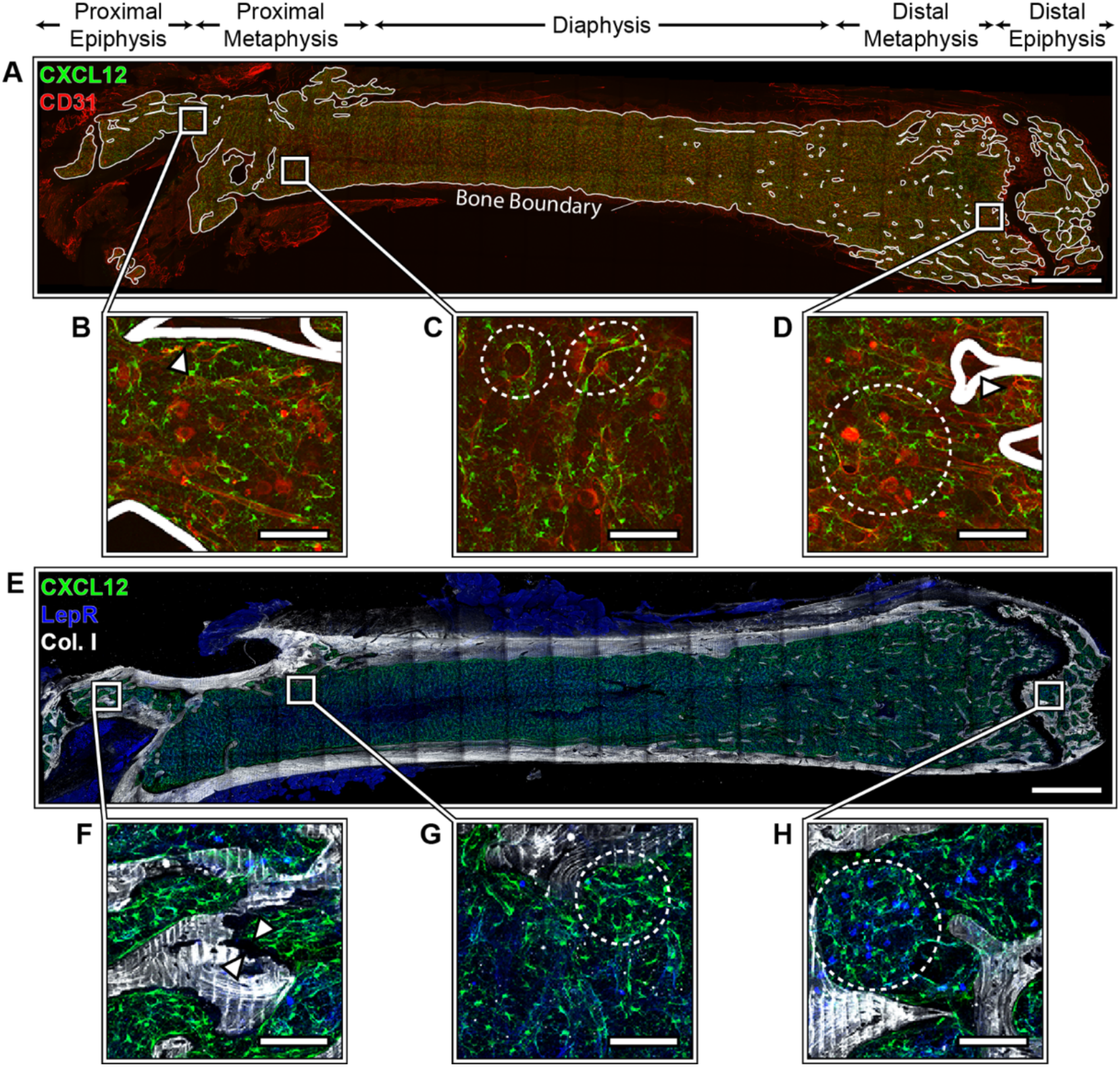
Additional Masked Immunofluorescent Images. Additional Immunofluorescent images of 12-week-old male mouse femurs used for pooled RSRK Analysis. Masks are shown in white. (A) CXCL12 (green) and CD31 (red), used in univariate CXCL12 analysis and bivariate CXCL12 vs. CD31 analysis. (B) CXCL12 (green), used in univariate CXCL12 analysis. (C) CXCL12 (green) and Collagen 1 (red) used in univariate CXCL12 analysis and bivariate CXCL12 vs. Bone analysis.

